# Ase1 domains dynamically slow anaphase spindle elongation and recruit Bim1 to the midzone

**DOI:** 10.1101/2020.07.20.211359

**Authors:** Ezekiel C. Thomas, Amber Ismael, Jeffrey K. Moore

## Abstract

How cells regulate microtubule crosslinking activity to control the rate and duration of spindle elongation during anaphase is poorly understood. In this study, we test the hypothesis that PRC1/Ase1 proteins use distinct microtubule-binding domains to control spindle elongation rate. Using budding-yeast Ase1, we identify unique contributions for the spectrin and carboxy-terminal domains during different phases of spindle elongation. We show that the spectrin domain uses conserved, basic residues to promote the recruitment of Ase1 to the midzone before anaphase onset and slow spindle elongation during early anaphase. In contrast, a partial Ase1 carboxy-terminal truncation fails to form a stable midzone in late anaphase, produces faster elongation rates after early anaphase, and exhibits frequent spindle collapses. We find that the carboxy-terminal domain interacts with the plus-end tracking protein EB1/Bim1 and recruits Bim1 to the midzone to maintain midzone length. Overall, our results suggest that the Ase1 domains provide cells with a modular system to tune midzone activity and control elongation rates.

## INTRODUCTION

The mitotic spindle is a complex microtubule network that divides labor between distinct subsets of microtubules. A fundamental question is how these subsets of microtubules are discretely regulated to perform specific roles within the spindle. The ‘midzone’ is a specific region within the spindle where interpolar microtubules that emanate from opposing halves of the spindle interdigitate to form a structure that stabilizes the spindle (McDonald et al., 1979; Winey et al., 1995). The midzone mechanically couples the two halves of the mitotic spindle as it undergoes major rearrangements in microtubule organization during spindle assembly and anaphase.

During anaphase B, the spindle elongates to move the two poles and associated genomes to separate regions of the dividing cell. Spindle elongation is a balancing act that involves binding and crosslinking antiparallel microtubules within the midzone and maintaining those interactions while the interpolar microtubules slide past each other to push apart the spindle poles. Disruption of midzone organization impairs spindle elongation and interferes with cytokinesis (Pellman et al., 1995; Jiang et al., 1998; Sharp et al., 1999a; Mollinari et al., 2002). The continued elongation of the spindle during anaphase B involves the coordination between the sliding of crosslinked microtubules within the midzone and the regulation of interpolar microtubules dynamics. Microtubules within the spindle are generally highly dynamic, but interpolar microtubules must be stabilized and grow to support the continued elongation of the spindle (Brust-Mascher et al., 2004; Cheerambathur et al., 2007; Fridman et al., 2009; Higuchi and Uhlmann, 2005; Maddox et al., 2000; Mallavarapu et al., 1999). The organization of the midzone influences the dynamics of the crosslinked microtubules (Bratman and Chang, 2007; Fridman et al., 2009), suggesting this organization is important for regulating interpolar microtubule dynamics.

Multiple conserved microtubule-associated proteins contribute to the organization of the midzone. Molecular motors of the kinesin-14 and kinesin-5 families crosslink microtubules and generate forces along interpolar microtubules that contribute to spindle elongation, making them a key component of the spindle. During anaphase, kinesin-5 driven plus-end motility within the midzone generates outward forces that elongate the spindle (Hoyt et al., 1992; Saunders and Hoyt, 1992; Sharp et al., 1999b, 2000; Straight et al., 1998), while kinesin-14s generate an inward force and provide structural support through microtubule crosslinking (Cai et al., 2009; Gardner et al., 2008; Saunders and Hoyt, 1992; Sharp et al., 1999a). Complete loss of kinesin-5 results in inviable cells due to spindle instability (Roof et al., 1992; Saunders and Hoyt, 1992); however, the simultaneous depletion of both kinesin-5 and kinesin-14 restores spindle viability (Rincon et al., 2017; Saunders and Hoyt, 1992). These results suggest that while molecular motors play key roles within the midzone, they are not essential for midzone function.

When cells are depleted for kinesin-5 and kinesin-14 motors, midzone function depends on the MAP65 family of non-motor, antiparallel microtubule crosslinkers (Rincon et al., 2017). The founding member of this family, MAP65, was discovered as a microtubule bundler in plants (Chang-Jie and Sonobe, 1993). Subsequently, homologues in budding yeast (Ase1) and humans (PRC1) were identified and recognized as important regulators of anaphase and cytokinesis (Jiang et al., 1998; Pellman et al., 1995). All MAP65 family members preferentially crosslink antiparallel microtubules and organize the midzone, and some members may perform additional functions (Bieling et al., 2010; Janson et al., 2007; Loïodice et al., 2005; Schuyler et al., 2003; Subramanian et al., 2010). *In vitro* experiments with purified fission yeast Ase1 or human PRC1 demonstrate that when these proteins crosslink antiparallel microtubules, they resist microtubule sliding (Braun et al., 2011; Gaska et al., 2020; Lansky et al., 2015). Consistent with these results, loss of ase1p or PRC1 in cells results in faster rates of spindle elongation (Janson et al., 2007; Pamula et al., 2019). How MAP65 family members are dynamically regulated in cells to set the timing, speed and magnitude of spindle elongation remains unclear. Furthermore, while crosslinking activity is relatively well studied, we have a comparatively poor understanding of how crosslinking activity is coordinated with the regulation of interpolar microtubule dynamics during anaphase B.

In this study, we sought to address these questions by investigating Ase1 function during spindle elongation in the budding yeast model, which offers precise genetic manipulation, control of expression levels, and a simplified mitotic spindle. The mitotic spindle in budding yeast consists of approximately 40 microtubules, the minus ends of which are stably anchored to two spindle pole bodies (SPBs) (Maddox et al., 2000; Winey et al., 1995). Thirty-two of these microtubules attach to the kinetochores of the 16 sister chromatids, and the remaining 8 are interpolar microtubules that interdigitate to form the midzone (O’Toole et al., 1999; Winey et al., 1995). In anaphase B, budding yeast Ase1 and homologues of kinesin-5 and kinesin-14 elongate the spindle to ~5X its metaphase length, delivering a copy of the duplicated genome to the bud and retaining the other copy in the mother (Winey and Bloom, 2012). During elongation, the number of interpolar microtubules decreases from 8 to approximately 2; at the completion of mitosis, the midzone is disassembled and interpolar microtubules depolymerize (Maddox et al., 2000; Winey and Bloom, 2012; Winey et al., 1995; Woodruff et al., 2010).

To understand the role of Ase1 in these different stages of spindle function, we used information from previous structural studies to generate mutations in chromosomal Ase1 and disable specific protein functions. We find that distinct domains of Ase1 are required for different stages of anaphase spindle elongation, and we identify a novel role for Ase1 in recruiting the end-binding (EB) protein homologue Bim1 to the spindle midzone. This recruitment is important for maintaining the size of the midzone during anaphase and stabilizing interpolar microtubules to sustain spindle elongation. Our findings demonstrate a modular complexity built into Ase1 that allows it to tune midzone activity and precisely regulate spindle elongation.

## RESULTS

### Ase1 domains uniquely contribute to slowing anaphase spindle elongation rates

MAP65 family members possess a stereotypic domain architecture: a structured amino-terminal domain, a central spectrin domain that binds microtubules, and an unstructured, intrinsically disordered carboxy-terminal domain (Kapitein et al., 2008; Kellogg et al., 2016; Mollinari et al., 2002; Schuyler et al., 2003; Subramanian et al., 2010). The amino-terminal domain supports homodimerization of Ase1/ PRC1 and is necessary for microtubule crosslinking activity (Janson et al., 2007; Schuyler et al., 2003; Subramanian et al., 2013). The spectrin domain interacts with a specific binding site on the microtubule surface, near the primary binding site for kinesin and dynein (Kellogg et al., 2016; Subramanian et al., 2010). The unstructured carboxy-terminal tail is necessary for microtubule binding, most likely through electrostatic interactions (Kapitein et al., 2008; Kellogg et al., 2016; Mollinari et al., 2002; Subramanian et al., 2010).

Based on these previous studies, we hypothesized that interactions between Ase1 and microtubules are primarily mediated by the spectrin and carboxy-terminal domains (Supplemental Figure 1). We disrupted the spectrin domain with three alanine substitution mutations R574A, K580A, and R581A (ase1^3A^; Figure 1A), which correspond to conserved basic amino acids in PRC1 that are predicted to bind to the acidic carboxy-terminal tail of β-tubulin (Figure 1A) (Kellogg et al., 2016). We disrupted the carboxy-terminal domain of Ase1 by generating truncated alleles. Removing the entire carboxy-terminal domain of Ase1 prevents localization to the nucleus (ase1^650Δ^; Supplemental Figure 2). We therefore generated a partial carboxy-terminal truncation that exhibits nuclear localization that is similar to full length Ase1 (ase1^Δ693^; Figure 1A; Supplemental Figure 2). All alleles were generated at the native chromosomal locus to maintain expression levels and provide the only source of Ase1 protein in the cell, and each allele is tagged with GFP at the carboxy-terminus. We predicted that these mutations would weaken the interaction between Ase1 and microtubules and result in faster anaphase elongation rates.

**Figure 1.**
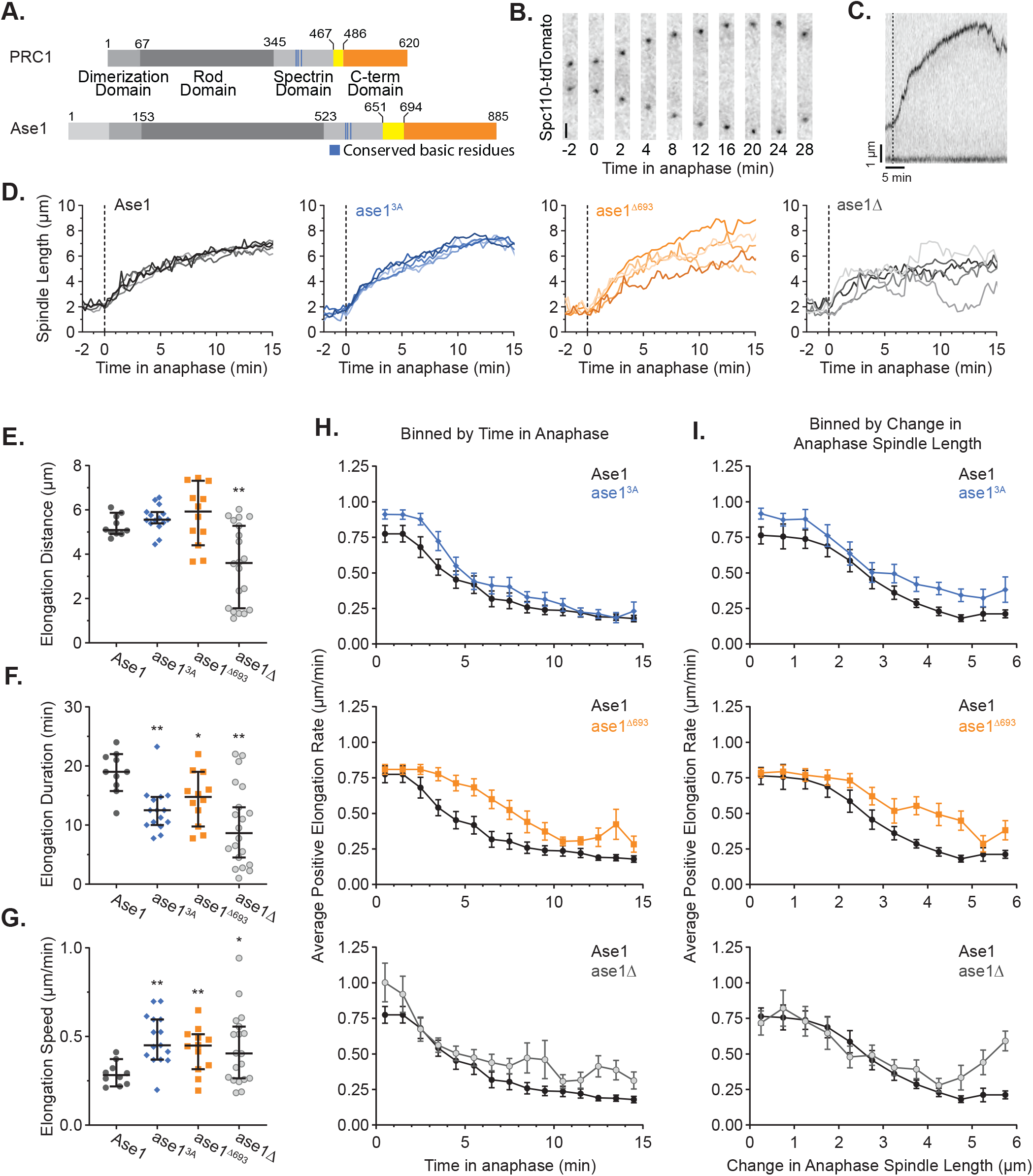
Ase1 Domains Uniquely Contribute to Anaphase Elongation Rates. A) Domain architecture of PRC1 and Ase1. B)Time lapse images of SPBs labeled with Spc110-tdTomato in a wild-type cell at 30°C. Scale bar = 1 μm. C) Time lapse imaging of cell in (B) shown as a kymograph aligned to one of the SPBs. Vertical dotted line represents the start of anaphase. D) Spindle length versus time in anaphase shown for five example cells for each genotype. Spindle length was determined by the three-dimensional distance between the SPBs. E) The change in spindle length (ase1^3A^, p = 0.135; ase1^Δ693^, p = 0.298; ase1Δ, p < 0.001), F) duration of elongation (ase1^3A^, p < 0.001; ase1^Δ693^, p = 0.016; ase1Δ, p < 0.001), and G) speed of elongation during anaphase spindle elongation events (ase1^3A^, p < 0.001; ase1^Δ693^, p = 0.005; ase1Δ, p = 0.016). Bars indicate median ± 95% CI. H) Average rates from elongation events binned by one-minute increments from anaphase onset. Data from wild-type controls is shown in black in each graph for comparison. Error bars represent 95% CI of the mean. I) Average rates from elongation events binned by half-micron increments of change in spindle length from anaphase onset. Data from wild-type controls is shown in black in each graph for comparison. Error bars represent 95% CI of the mean. For all panels, a minimum of ten cells were analyzed for each genotype. *p < 0.05, **p < 0.01, as determined by t-test with Welch’s correction compared to wild type.

To test this prediction and determine the contributions of Ase1’s spectrin and carboxy-terminal domains to elongation rates, we used spinning disc confocal microscopy to measure spindle elongation over time in living cells (Figure 1B and C, Material and Methods). We collected Z-series at 15-second intervals and defined spindle elongation as an increase in the three-dimensional distance between the SPBs. While wild-type cells display relatively consistent spindle elongation rates, *ase1*^3A^, *ase1*^Δ693^, and *ase1*Δ null mutant cells exhibit spindle elongation rates that are variable from cell-to-cell of the same genotype, or even within the anaphase of a single cell (Figure 1D). To account for this variability, we developed a linear-regression based model to identify “elongation events”, defined as periods of spindle elongation lasting at least 60 seconds (5 time points, Materials and methods), and analyzed the spindle length changes and durations of these periods. For wild-type cells these events appeared to encompass all of anaphase with a median spindle length increase of 5.09 μm [4.92, 5.87] (median [(95% CI lower limit, 95% CI upper limit], Figure 1E). The median duration of these events was 19.0 min [15.8, 22.0] (Figure 1F), resulting in a median elongation speed of 0.28 μm/min [0.22, 0.37] (Figure 1G). Compared to wild-type cells, spindle elongation in *ase1*Δ cells is frequently interrupted by periods where spindles shorten (Figure 1D). Accordingly, elongation events in *ase1*Δ cell result in significantly smaller length changes and are short-lived, with a median of 3.61 μm [1.57, 5.28] and 8.63 min [4.50, 13.0] (p < 0.001 t-test with Welch’s correction, Figure 1E and F). However, the elongation speed in *ase1*Δ cells is significantly faster than that of wild-type cells (median = 0.41 μm/min [0.27, 0.56], p = 0.016, t-test with Welch’s correction, Figure 1G). These results are consistent with the role of Ase1 in slowing anaphase spindle elongation.

Mutations disrupting the spectrin or carboxy-terminal domains disturb Ase1’s role in slowing anaphase spindle elongation. Both *ase1*^3A^ and *ase1*^Δ693^ mutant cells display changes in spindle length similar to that of wild-type cells (*ase1*^3A^ = 5.56 μm [5.39, 5.91]; *ase1*^Δ693^ = 5.93 μm [4.41, 7.32]); however, the median duration of these events is reduced (ase1^3A^ = 12.5 min [10.0, 14.8]; ase1^Δ693^ = 14.8 min [9.75, 19.0]). Thus, the elongation speeds are significantly faster in these mutants (*ase1*^3A^ = 0.45 μm/min [0.37, 0.60], p < 0.001; *ase1*^Δ693^ = 0.45 μm/min [0.32, 0.51], p < 0.01; t-test with Welch’s correction, Figure 1E-G). These results suggest that the spectrin and carboxy-terminal domains of Ase1 regulate the speed of anaphase spindle elongation.

To precisely characterize how these domains contribute to spindle elongation, we generated composite profiles of spindle elongation by averaging intervals of spindle elongation lasting at least 1 minute from populations of cells (see Materials and Methods, Supplemental Figure 3, Supplemental Table 1). Consistent with previous studies(Kahana et al., 1995; Straight et al., 1997, 1998), our results demonstrate that cells exhibit a fast rate of spindle elongation upon entering anaphase, and that rate slows as cells progress through anaphase and the spindle nears its terminal length. For wild-type cells, an initial elongation rate of 0.77 μm/min is sustained for approximately 2 minutes (Figure 1H), or when the spindle has elongated by 1.5 μm (Figure 1I), and then gradually decreases to a slower rate of approximately 0.20 μm/min. Spindle elongation in *ase1Δ* null mutants differs in three respects: 1) the initial elongation rate is faster (1.0 μm/min), 2) the subsequent rate decrease is more pronounced, and 3) rates during the later phase are highly variable and tend to be faster (Figure 1H and I).

Mutations disrupting the spectrin or carboxy-terminal domains of Ase1 elicit specific phenotypes in this analysis. Cells expressing *ase1*^3A^ exhibit faster initial elongation rates than wild-type controls, followed by a steady decrease that is reminiscent of wild type (Figure 1H). Figure 1I shows that the *ase1*^3A^ cells begin to slow after elongating the spindle by 1.5 μm, which is similar to the transition point observed in wild-type cells. These results suggest that the spectrin-homology domain of Ase1 normally slows spindle elongation during early anaphase. In contrast, *ase1*^Δ693^ mutant cells have initial elongation rates that are similar to those of wild-type cells but sustain faster elongation rates further into anaphase and for longer spindle lengths (Figure 1 H and I). These results suggest that the carboxy-terminal domain of Ase1 normally slows elongation in late anaphase, as the spindle approaches its terminal length. Together, our results indicate distinct roles for Ase1 domains in regulating elongation as the spindle transitions to longer lengths.

### Ase1 spectrin and carboxy-terminal domains stabilize the spindle

In addition to regulating the rate of elongation, crosslinking by Ase1 is important for supporting spindle structure as it reaches longer lengths. Consistent with this, we observed greater spindle length fluctuations over time in *ase1* mutant cells. Whereas wild-type cells exhibit sustained spindle elongation during anaphase, *ase1* mutant cells exhibit frequent interruptions by sharp increases or decreases in spindle length (Figure 2A). When comparing populations of cells, this manifests as an increased standard deviation of spindle length and is more pronounced later in anaphase (Figure 2B). To quantify this variability in spindle length across populations, we compared the standard deviation of spindle length over time aligned to the onset of anaphase for at least 10 cells of each genotype (Figure 2C). In this analysis, wild-type cells have a flat standard deviation indicating that spindle elongation during anaphase is consistent across cells, while *ase1*Δ null cells exhibit significantly more biological variation throughout anaphase (p < 0.05 F-test, Supplemental Table 2). Cells expressing *ase1*^*3A*^ resemble wild-type cells for the first 3.5 minutes, after which there are periods of greater variation. Cells expressing *ase1*^*Δ693*^ diverge from wild-type cells after 3 minutes and continue to exhibit increased variation through late anaphase.

**Figure 2.**
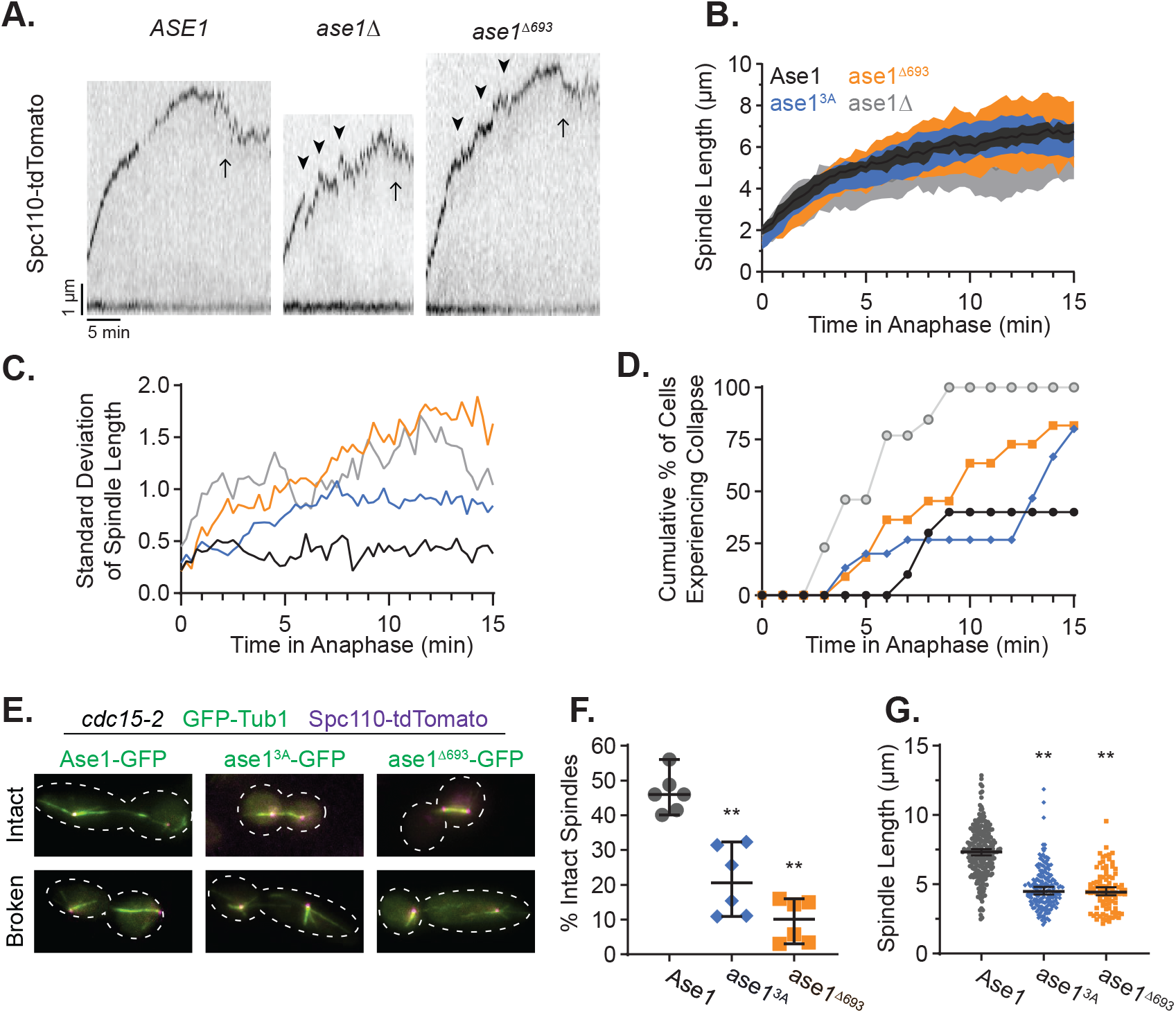
Ase1 Domains Stabilize the Spindle. A) Example kymographs of *ASE1*, *ase1Δ* and *ase1*^Δ693^ cells aligned to one SPB. Arrows represent spindle disassembly at the end of mitosis. Arrowheads represent collapses during anaphase spindle elongation. B) Average spindle length over time as measured by the three-dimensional distance between the SPBs. Shaded region represents standard deviation. A minimum of ten cells were measured for each genotype. C) The standard deviation of spindle length over time for the same cells in (B). D) The cumulative percent of cells that have experienced a spindle collapse event, as a function of time in anaphase. E) Example images of fixed *cdc15-2* cells arrested at 37°C for 3 hours. Cells were considered to have intact or broken spindles as determined by continuity of GFP-tubulin fluorescence. GFP-Tub1 (green); Ase1-GFP or ase1^3A^-GFP or ase1^Δ693^-GFP (green); Spc110-tdTomato (magenta). Scale bar = 1 μm. F) Percentage of *cdc15-2* arrested cells with intact spindles. Each data point represents a minimum of one-hundred cells from a separate experiment. Bars are median ± 95% CI. G) Spindle lengths from cells in (F) with intact spindles. Bars are median ± 95% CI. **p < 0.01, as determined by t-test for (F) and Mann-Whitney *U* test in (G) compared to wild type.

We reasoned that the increased variability in spindle length may reflect defects in spindle stability. To test this, we quantified spindle collapse events during anaphase, defined as a decrease in spindle length of at least 0.5 um that occurs within a one-minute period (5 time points; Figure 2A, see arrowheads). These collapse events are likely to be distinct from spindle disassembly at mitotic exit because they occur within 15 minutes of anaphase onset and spindles typically recover from a collapse event and resume elongation. Our analysis revealed that spindle collapses are more common in *ase1* mutants than in wild-type controls. For *ase1*Δ null mutants, spindle collapses are evident in early anaphase and all cells exhibit collapses within 9 minutes of entering anaphase (Figure 2D). *ase1*^693Δ^ mutant cells start to exhibit spindle collapses after several minutes in anaphase, and these collapses accumulate during late anaphase (Figure 2D). The phenotype of *ase1*^693Δ^ mutant cells is more severe than that of *ase1*^3A^ mutants in this analysis. Notably, the trends of spindle collapse events over time is reminiscent of the trends for spindle length standard deviations for all *ase1* mutants (Figure 2C), consistent with the notion that variations in spindle lengths reflect spindle instability.

To test the possibility that spindle collapse events in *ase1* mutant cells could be attributable to early mitotic exit, we prevented mitotic exit and assayed the maintenance of intact spindles. We arrested cells in telophase using a temperature sensitive *cdc15-2* allele (Hartwell and Smith, 1985), fixed, and then imaged GFP-labeled microtubules to determine whether the spindle was still intact (Figure 2E). Mutants expressing *ase1*^Δ693^ rarely exhibit intact spindles during telophase arrest, while *ase1*^3A^ cells exhibit an intermediate level of intact spindles, compared to that of wild-type controls (p < 0.001, t-test; Figure 2F). Furthermore, when the spindle is intact in telophase-arrested cells, the median spindle length is significantly lower for *ase1*^3A^ (median = 4.49 μm [4.27, 4.81]) and *ase1*^Δ693^ (median = 4.44 μm [4.20, 4.78]) mutant cells, relative to that of wild-type cells (median = 7.35 μm [7.09, 7.54]; p < 0.001, Mann-Whitney U test; Figure 2G). Altogether, these results suggest that both the spectrin domain and the carboxy-terminal domain of Ase1 stabilize the anaphase spindle, but the carboxy-terminal domain is particularly important for spindle stability during late anaphase.

### Spectrin domain promotes Ase1 recruitment to the spindle

Next, we examined whether defects in anaphase spindle elongation and stability could be explained by reduced recruitment of ase1 mutant proteins to the mitotic spindle. To measure the recruitment of Ase1 to the spindle throughout mitosis, we collected single timepoint Z-series images of asynchronous cells and quantified the total intensity of Ase1-GFP signal along the spindle between the two SPBs, and compared total signal intensity as a function of spindle length (Figure 3A, Material and Methods). Wild-type Ase1-GFP displays a linear band of GFP signal between the two SPBs (Figure 3B), and the total intensity of this signal increases throughout mitosis (Figure 3C). To correlate Ase1 localization to different stages of spindle elongation, we divided images into three bins, representing pre-anaphase, the fast phase of anaphase spindle elongation, and the slow phase of anaphase spindle elongation (Kahana et al., 1995; Straight et al., 1997, 1998). These bins are defined for each genotype, based on the kinetics of spindle elongation measured in Figure 1, and are represented by vertical dotted lines in Figures 3C-E (Supplemental Figure 3, see Materials and Methods). The fluorescence intensity of wild-type Ase1 approximately doubles from pre-anaphase to anaphase but remains relatively constant during the fast and slow phases of anaphase (Figure 3F-H).

**Figure 3.**
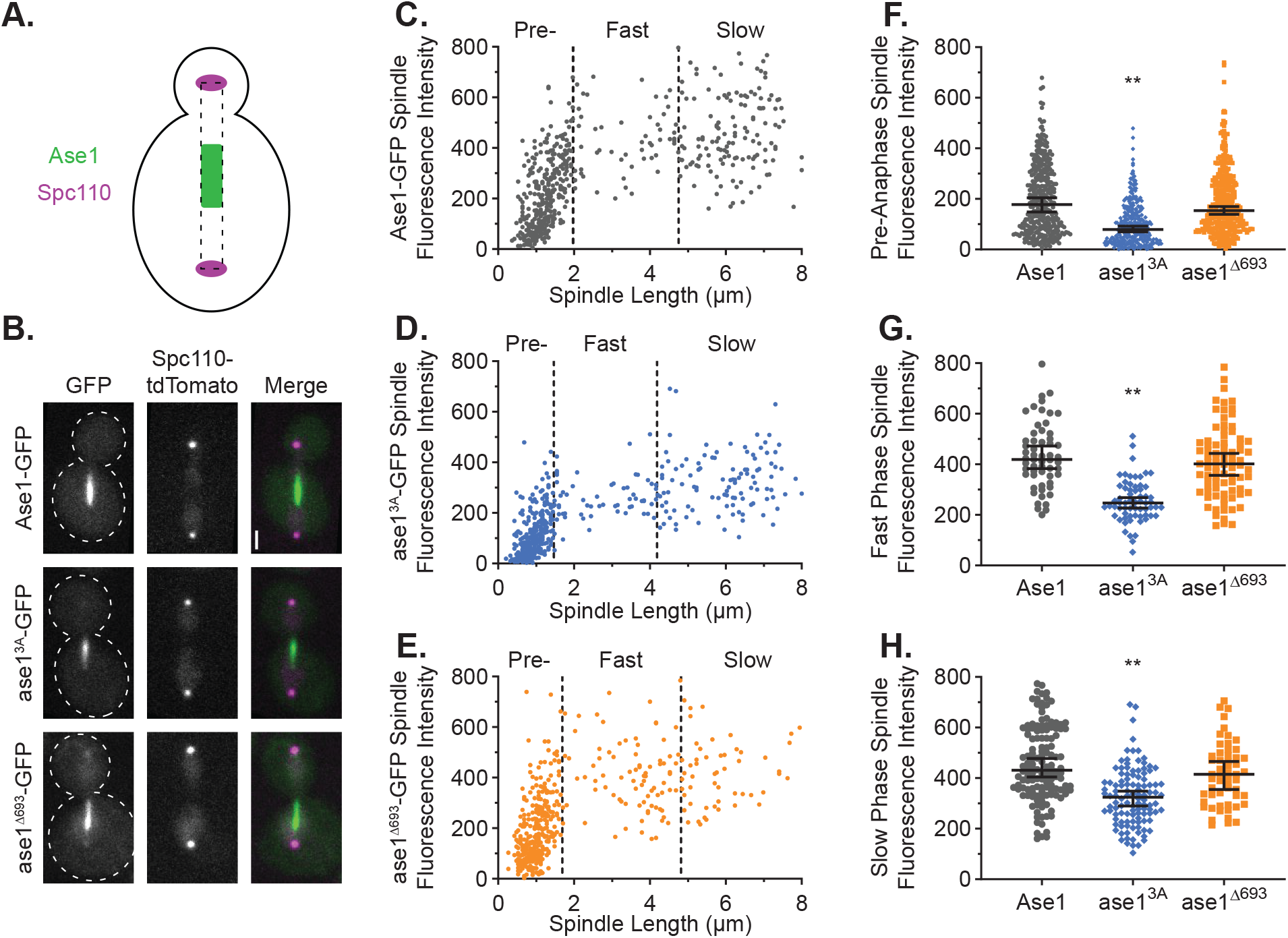
Spectrin Domain Promotes Ase1 Recruitment to the Spindle. A) Diagram of the analysis performed. Dotted box represents the area drawn along the spindle between the two SPBs and encompassing the Ase1-GFP region. B) Example images of cells in anaphase. Scale bar = 1 μm. C) Background-subtracted spindle fluorescence intensity for Ase1-GFP expressing cells as a function of spindle length, *n* = 370 cells; D) ase1^3A^-GFP, *n* = 360 cells; or E) ase1^Δ693^-GFP, *n* = 370 cells. Vertical dotted lines denote the determined boundaries between pre-anaphase, the fast phase of anaphase, and the slow phase of anaphase. F) GFP intensity values for pre-anaphase cells from (C-E) (ase1^3A^, p < 0.001; ase1^Δ693^, p = 0.173), G) for the fast phase of anaphase (ase1^3A^, p < 0.001; ase1^Δ693^, p = 0.351), or H) for the slow phase of anaphase (ase1^3A^, p < 0.001; ase1^Δ693^, p = 0.053). Bars are median ± 95% confidence interval. ** p < 0.01, as determined by Mann-Whitney *U* test for (F) and t-test with Welch’s correction for (G) and (H) compared to wild type.

Although ase1^3A^-GFP expressing cells display a linear band of GFP fluorescence between the two SPBs (Figure 3B), the level of signal is consistently lower at each stage of spindle elongation and does not reach the level of enrichment seen for wild-type Ase1-GFP (Figure 3D, F-H). Despite these decreased levels, we still observed an approximate doubling of ase1^3A^– GFP signal when comparing pre-anaphase cells to the fast phase of anaphase (Figure 3F and G). These results indicate that the spectrin domain promotes Ase1’s recruitment to the spindle prior to anaphase, and that conserved basic residues within this domain are particularly important.

Most cells expressing ase1^Δ693^-GFP display a linear band of GFP signal between the SPBs, with total signal intensities that are similar to those observed for wild-type Ase1-GFP (Figure 3B, E-H). This indicates that the carboxy-terminal domain is not required to recruit Ase1 to the spindle. However, we noticed that late anaphase cells expressing ase1^Δ693^-GFP exhibit higher signal in the nucleoplasm and an average of 26% of anaphase cells lack clear GFP enrichment on the spindle (Supplemental Figure 4; see below).

### Carboxy-terminal domain is necessary for stable retention of Ase1 at the spindle middle and organization of spindle microtubules

Although the total levels of ase1^Δ693^-GFP recruited to the spindle are similar to that of wild-type Ase1-GFP, we observed that the distribution of ase1^Δ693^-GFP is asymmetrically localized toward one end of the spindle (Figure 3B). We measured this asymmetry by analyzing the distribution of GFP signal along the spindle. We first identified the region of signal enrichment in anaphase cells, and then determined the position of the midpoint of this region relative to the geometric center of the spindle (Figure 4A, Materials and methods). For wild-type Ase1-GFP and ase1^3A^-GFP, the average midpoint is located at the center of the spindle throughout anaphase (Figure 4B and C). In contrast, ase1^Δ693^–GFP signal is shifted toward the mother cell, and this asymmetry becomes more pronounced over time as the spindle length increases during the transition from the fast to the slow phase of anaphase (Figure 4B and C). These results suggest that the carboxy-terminal domain is necessary for the retention of Ase1 at the center of the spindle.

**Figure 4.**
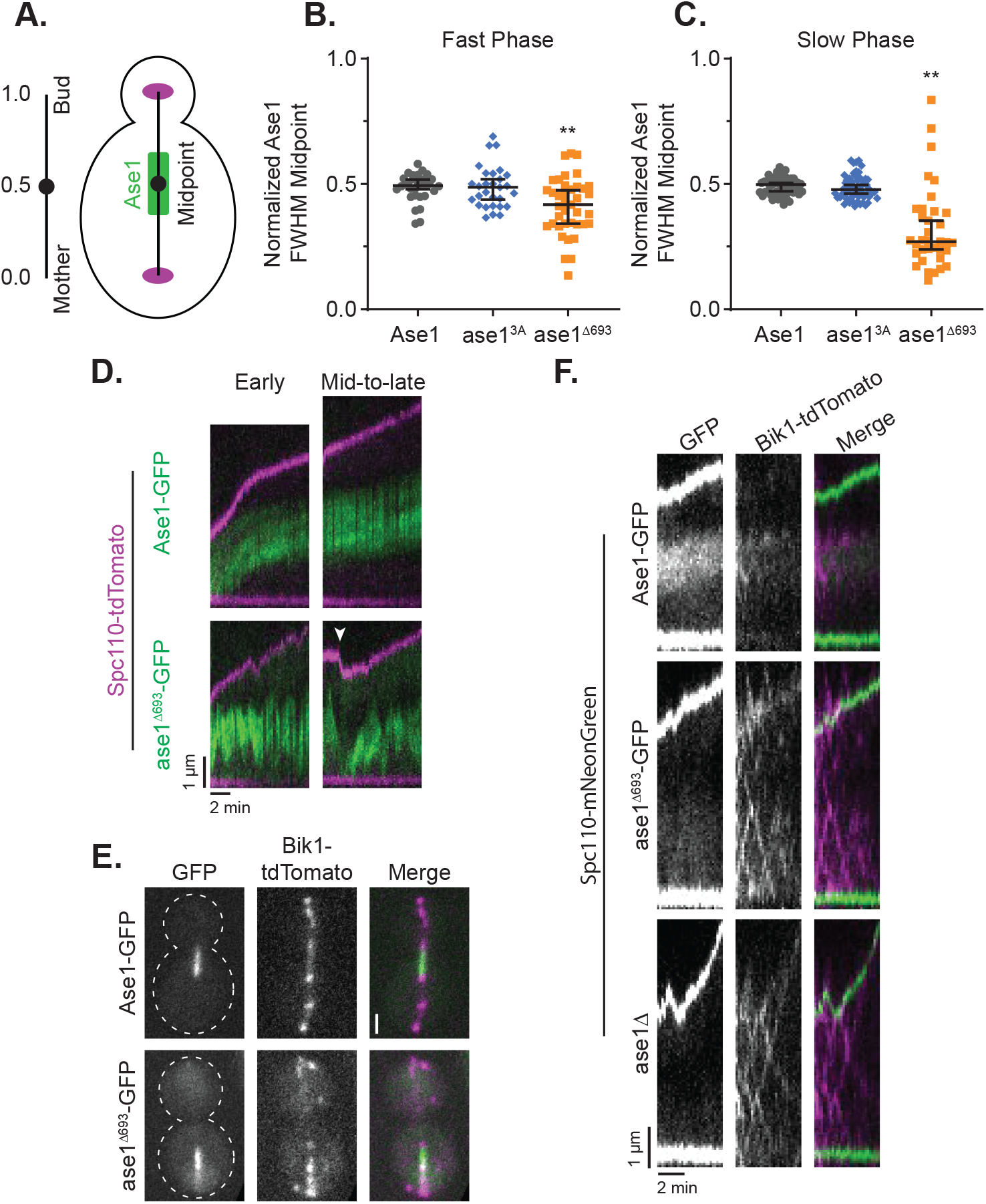
Carboxy-terminal Domain is Necessary for Spindle Microtubule Organization. A) Diagram of the analysis performed. Ase1 region was determined by full-width half-max analysis of GFP signal, and the geometric midpoint determined relative to the spindle length. A value of zero represents the SPB within the mother cell, a value of 0.5 is the geometric midpoint of the spindle, and a value of 1.0 is the SPB within the bud. B) Ase1 region midpoint during the fast phase of anaphase spindle elongation. Ase1-GFP, *n* = 31; ase1^3A^-GFP, p = 0.964, *n* = 29; ase1^Δ693^-GFP, p < 0.001, *n* = 39. C) Ase1 region midpoint during the slow phase of anaphase spindle elongation. Ase1-GFP, *n* = 52; ase1^3A^-GFP, p = 0.851, *n* = 56; ase1^Δ693^-GFP, p = 0.002, *n* = 35. D) Example kymographs of cells during early anaphase or mid-to-late anaphase showing SPBs labeled with Spc110-tdTomato (magenta) and Ase1-GFP (green). Arrowheads represent spindle collapse. Kymographs are aligned to the mother cell SPB along the bottom of each image. E) Example anaphase cells expressing Bik1-tdTomato and Ase1-GFP or ase1Δ693-GFP. Scale bar = 1 μm. F) Example kymographs of anaphase cells expressing Bik1-tdTomato and Ase1-GFP or ase1^Δ693^-GFP. Kymographs are aligned to the mother cell SPB along the bottom of each image. **p < 0.01, as determined by Mann-Whitney *U* test compared to wild type.

Given the variable localization of ase1^Δ693^-GFP observed in populations of cells, we next asked if the localization of ase1^Δ693^-GFP is dynamic within individual cells over time. To test this possibility, we collected z-series at 10-second intervals of cells starting at spindle lengths of less than 5 μm. From these time series we made kymographs in which we fixed the position of the SPB within the mother cell. As shown in Figure 4D, wild-type Ase1-GFP remains located at the center of the spindle as the spindle length increases (also see Supplemental Videos 1 and 2). In contrast, ase1^Δ693^–GFP displays a dynamic, asymmetric localization that not only shifts along the length of the spindle, but also can be transiently lost from the spindle (Figure 4D arrowhead, Supplemental Videos 3 and 4). In this example, loss of ase1^Δ693^–GFP from the spindle coincides with a spindle collapse and increase in nuclear GFP background, after which the ase1^Δ693^–GFP signal recovers and spindle elongation resumes (Figure 4D, Supplemental Video 4). We also observed short collapse events where ase1^Δ693^-GFP was not lost from the spindle (Supplemental Video 3), indicating that collapse does not require full loss of ase1^Δ693^-GFP. All together these results indicate that the carboxy-terminal domain promotes the stable retention of Ase1 at the center of the spindle, and that transient loss contributes to spindle collapse.

The asymmetric distribution of ase1^Δ693^ on the spindle could be explained by two competing hypotheses: 1) the carboxy-terminal domain is necessary for the preferential binding of Ase1 to antiparallel microtubules or 2) the carboxy-terminal domain is necessary for the retention of antiparallel microtubules at the spindle center. To test these hypotheses, we imaged CLIP170/Bik1 to determine the location of microtubule plus ends in the anaphase spindle. As expected, Bik1-tdTomato foci localize to the ends of the wild-type Ase1-GFP region at the center of the spindle (Figure 4E). In cells expressing ase1^Δ693^ the Bik1-tdTomato foci localize to the ends of the asymmetric ase1^Δ693^-GFP region, suggesting the entire midzone is asymmetrically localized along the spindle. We imaged the microtubule plus ends by time-lapse microscopy and confirmed that Bik1 remains associated with the wild-type Ase1-GFP region as the spindle elongates (Figure 4F). In cells expressing ase1^Δ693^, Bik1 foci are highly dynamic and travel along the length of the spindle. We also observe moving Bik1 foci in *ase1*Δ null cells (Figure 4F). Although we cannot determine which SPB these microtubules emanate from, these results suggest that Ase1 normally contributes to stabilizing the plus ends of interpolar microtubules at the center of the spindle, and that this requires the carboxy-terminal domain.

### Carboxy-terminal domain recruits Bim1 to the midzone through SxIP motif

To determine how the carboxy-terminal domain stabilizes plus ends, we tested for interactions with a panel of mitotic spindle proteins. Through a two-hybrid assay we identified a positive interaction between full-length Ase1 and EB1/Bim1, a microtubule plus end tracking protein (Figure 5A). The partial carboxy-terminal domain of Ase1 (residues 694-885) was necessary and sufficient for this interaction, suggesting that this domain interacts with Bim1 (Figure 5A). A wide range of proteins interact with the EB1/Bim1 proteins through SxIP motifs (Honnappa et al., 2009; Jiang et al., 2012), and we identified an SxIP motif within the Ase1 carboxy-terminal domain (SHIP, residues 775-8). To test the relevance of this motif in the Ase1-Bim1 interaction, we generated mutations predicted to disrupt binding to Bim1 (I777N and P778N). Consistent with our prediction, altering the SHIP motif ablated Ase1’s two-hybrid interaction with Bim1 (Figure 5A). These results suggest that the Ase1 carboxy-terminal tail domain interacts with Bim1 through a canonical SxIP motif.

**Figure 5.**
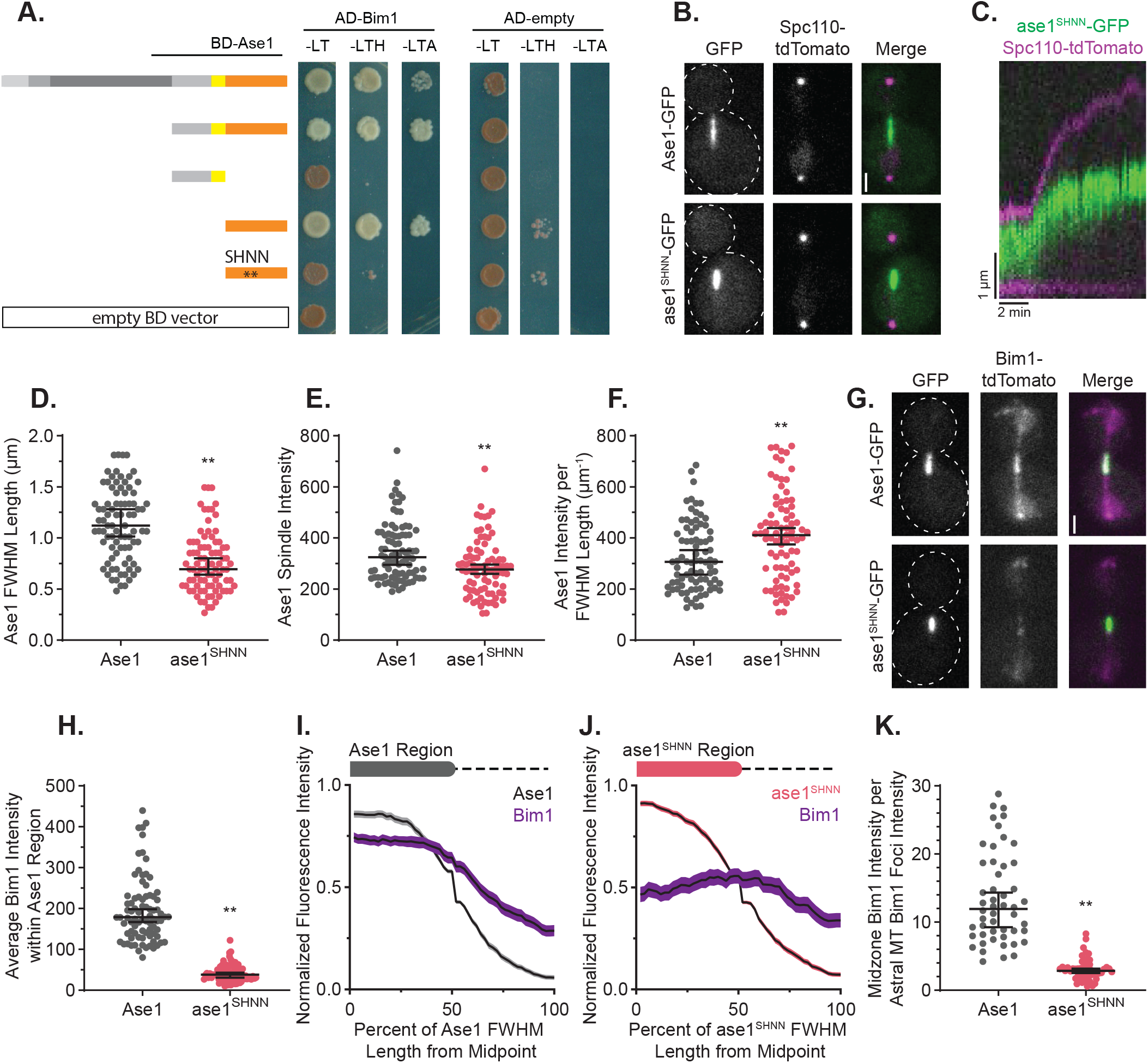
Ase1 Recruits Bim1 to the Midzone. A) Two-hybrid assay for Ase1 interactions. Cells transformed with Gal4-DNA binding domain fused to full length or truncations of Ase1 (rows) and either Gal4-activation domain fused to Bim1 or Gal4-activation domain alone (AD-empty) were plated onto media to select for plasmid retention (-LT, -Leucine -Tryptophan), or selecting for plasmid and interaction (-LTH, -Leucine - Tryptophan -Histidine; -LTA, -Leucine -Tryptophan -Adenine) and incubated at 30°C for 2 days. The two asterisks represent two point mutations within the Ase1 carboxy-terminal domain (I777N and P778N). B) Example images of cells expressing Ase1-GFP or ase1^SHNN^-GFP and Spc110-tdTomato. Scale bar = 1 μm. C) Example kymograph of a cell expressing ase1^SHNN^-GFP (green) and Spc110-tdTomato (magenta) progressing through early anaphase. Kymograph is aligned to the mother cell SPB along the bottom of the image. D) Length of the Ase1-GFP (n = 85 cells) or ase1SHNN-GFP (n = 83 cells) region as determined by full-width half-max analysis for cells with spindle lengths greater than 4 μm (p < 0.001). Bars are median ± 95% CI. E) Background-subtracted total spindle fluorescence intensity of Ase1-GFP or ase1^SHNN^-GFP for cells with spindle lengths greater than 4 μm (p < 0.001). Bars are median ± 95% confidence interval. F) Ase1-GFP or ase1^SHNN^-GFP fluorescence intensity per length of the GFP region for each of the cells in (D) and (E) (p = 0.004). Bars are median ± 95% CI. G) Example images of cells expressing Bim1-tdTomato (magenta) and Ase1-GFP or ase1SHNN-GFP (green). Scale bar = 1 μm. H) Average Bim1-tdTomato fluorescence intensity along a line scan corresponding to the Ase1-GFP (n = 86 cells) or ase1SHNN-GFP (n = 88 cells) full-width half-max region (p < 0.001). Bars are median ± 95% CI. I) Normalized Bim1-tdTomato and GFP fluorescence intensity relative to the Ase1-GFP or J) ase1^SHNN^-GFP region. Each spindle was analyzed from the midpoint of the Ase1 region (defined as 0 on the x-axis) as determined by full-width half-max. 50 on the x-axis represents the FWHM of the half of the Ase1 region, and 100 represents 1X distance beyond the Ase1 FWHM. These regions are illustrated by a diagram along the top of each graph. Lines are average values with shaded regions representing ± 95% CI. Ase1, n = 86; ase1^SHNN^, n = 88. K) Ratio of midzone Bim1-tdTomato fluorescence over intensity at a single microtubule plus end. Total Bim1-tdTomato signal at the Ase1-GFP (n = 50 cells) or ase1^SHNN^-GFP (n = 51 cells) defined midzone was divided by the total Bim1-tdTomato at a single astral microtubule plus end within the same cell (p < 0.001). Bars are median ± 95% CI. **p < 0.01, determined by Mann–Whitney *U* test.

We next tested the role of Bim1 binding in Ase1 function. We generated a full-length, GFP tagged *ase1* allele at its native locus with point substitutions I777N and P778N to disrupt interactions with Bim1 (ase1^SHNN^; Figure 5B). ase1^SHNN^-GFP localizes to a region at the middle of the spindle (Figure 5B and C, quantification not shown), suggesting the interaction with Bim1 is not necessary to stabilize the midzone at the center of the spindle. However, the region of ase1^SHNN^-GFP signal was shorter than that of wild-type Ase1-GFP, implying a shorter midzone (Figure 5D). The amount of ase1^SHNN^-GFP on the spindle was slightly but statistically significantly less than that of wild-type Ase1-GFP (Figure 5E). When adjusted for the reduction in midzone length, the amount of ase1^SHNN^-GFP intensity per midzone length is greater than that of wild-type Ase1-GFP (Figure 5F). These results suggest that interaction with Bim1 is necessary to prevent the compaction of the midzone.

Bim1 does not localize to the midzone in *ase1*Δ null cells (Khmelinskii et al., 2007), but this could be attributed to lack of recruitment by Ase1 or lack of the midzone organization itself. We next imaged cells expressing Bim1-tdTomato to test the prediction that Bim1 is recruited to the midzone by Ase1’s carboxy-terminal domain. While Bim1 colocalizes with wild-type Ase1 at the midzone, cells expressing ase1^SHNN^ exhibit significant reduction of Bim1 recruitment to the midzone (Figure 5G and H). Bim1 is expected to directly bind to the plus ends of interpolar microtubules, so we examined the distribution of Bim1 on the spindle relative to the Ase1 region. In cells expressing wild-type Ase1-GFP, Bim1-tdTomato levels are evenly distributed throughout the Ase1 region and begin to decrease at the ends (Figure 5I). In cells expressing ase1^SHNN^-GFP, Bim1-tdTomato is slightly enriched at the ends of the ase1^SHNN^ region, where the microtubule plus ends are predicted to be located, but not enriched throughout the ase1^SHNN^ region (Figure 5J). These results suggest that when interaction with Ase1 is disrupted, a smaller amount of Bim1 still associates with the plus ends of interpolar microtubules.

Finally, we asked whether interaction with Ase1 recruits additional Bim1 to the midzone, beyond the amount that would be expected to be recruited through microtubule plus-end binding alone. To compare the amount of Bim1-tdTomato signal at the anaphase midzone to that at an individual microtubule plus end, we divided the total amount of Bim1-tdTomato within the midzone by the amount of Bim1 found at the plus end of a single astral microtubule in the same cell. This analysis shows that the median amount of Bim1 present at a wild-type midzone is equivalent to the cumulative amount of 12 plus ends (Figure 5K). Electron tomography of budding yeast cells shows that as the spindle elongates during anaphase, the number of interpolar microtubules decreases from approximately 8 to 2 (Winey and Bloom, 2012; Winey et al., 1995). Therefore, the amount of Bim1 present at the midzone in wild-type cells is at least 50% greater than would be expected from plus end binding alone. In contrast, the median amount of Bim1 present at the midzone of cells expressing ase1^SHNN^ is equivalent to 3 plus ends (Figure 5K). This suggests that disrupting the Ase1-Bim1 interaction not only depletes Bim1 from the midzone, but it may also decrease the number of interpolar microtubule plus ends in the midzone. All together these results indicate that Ase1 enriches Bim1 at the midzone through an SxIP motif within the carboxy-terminal domain of Ase1.

## DISCUSSION

In this study, we show that the spectrin and carboxy-terminal domains of Ase1 regulate distinct aspects of spindle elongation during anaphase, and that these activities involve distinct interaction modes with microtubules and the plus-end binding protein Bim1/EB1. Furthermore, our findings give new insight into an underappreciated role of the midzone in regulating microtubule dynamics to stabilize the spindle as it reaches long lengths. Together, our results demonstrate that Ase1 is more than a microtubule crosslinking protein; it is a modular hub for coordinating local control of dynamic instability amongst a subset of microtubules in the spindle.

Our results indicate that Ase1 uses its spectrin domain for recruitment to the midzone and for setting the rate of anaphase spindle elongation. Spectrin domains are found in a wide variety of proteins (e.g. α-actinin, dystrophin, spectraplakins, and nesprins) that act as bridges between cytoskeletal networks or between these networks and membrane-bound proteins. In these examples, the spectrin domains commonly act as flexible spacers that fold and unfold in response to mechanical forces and in some cases interact with signaling proteins or cytoskeletal regulators (Chen et al., 2006; Liem, 2016; Noordstra et al., 2016; Sonnenberg et al., 2007). The spectrin domain of Ase1 is unique in that it provides direct microtubule-binding activity that is conserved across species. Previous studies found that protein truncations containing the spectrin domains of fission yeast Ase1, Arabidopsis MAP65-1, and mammalian PRC1 all exhibit direct microtubule-binding activity (Kapitein et al., 2008; Li et al., 2007; Mollinari et al., 2002; Smertenko et al., 2004; Subramanian et al., 2010). Structural studies of PRC1 identified positively charged amino acids in helix 8 of the spectrin domain that increase PRC1 affinity for microtubules through electrostatic interactions (Kellogg et al., 2016; Subramanian et al., 2010). Ablating the charges of these residues by replacing them with alanines reduces the affinity of PRC1 for microtubules and increases the distance between segregated chromosomes in anaphase (Kellogg et al., 2016; Pamula et al., 2019). Our work extends these findings by identifying the role of these positively charged amino acids in mitotic spindle elongation. We find that replacing the analogous amino acids in budding yeast Ase1 with alanines increases spindle elongation rates during anaphase (Figure 1) and diminishes localization to the spindle (Figure 3). We conclude that these positively-charged residues are a conserved feature of Ase1’s spectrin domain that promotes recruitment to the spindle before anaphase onset and slows spindle elongation during early anaphase.

The faster rate of spindle elongation during early anaphase in the ase1^3A^ mutant is likely attributable to the diminished recruitment of the ase1^3A^ protein to the midzone. Previous *in vitro* studies with purified fission yeast Ase1 established that the amount of ase1p that is recruited to a region of antiparallel microtubule overlap is proportional to the magnitude of braking force that resists the plus-end directed sliding of those microtubules (Braun et al., 2011; Lansky et al., 2015). Furthermore, a recent study found that the braking activity of PRC1 is proportional to the amount of PRC1 that is recruited to microtubule overlaps and is highly sensitive to ionic strength (Gaska et al., 2020). We propose that electrostatic interactions between the positively charged amino acids in the Ase1/PRC1 spectrin domain and the negatively charged microtubule surface promote recruitment to the midzone and microtubule crosslinking activity. This model of electrostatic interactions is appealing because it invites potential regulation of braking activity through posttranslational modifications at the level of either Ase1 or the microtubule surface (see below). Together, these could provide cells with a two-dimensional mechanism for fine tuning crosslinking activity and the rate of anaphase spindle elongation.

The unstructured carboxy-terminal domain of Ase1 appears to support several functions in cells. The proximal region of the carboxy-terminal domain that is immediately adjacent to the spectrin domain is required for the localization of Ase1 to the nucleus (residues 651-693; Supplemental Figure 2). The requirement for nuclear localization, plus the presence of multiple lysine or arginine residues, suggests that this region may be involved in binding importins and acting as a nuclear localization signal for Ase1 (Supplemental Figure 1). A recent study of PRC1 found that a truncation of the carboxy-terminal domain that removed the region analogous to 653-693 completely ablated localization to the spindle (Pamula et al., 2019). Together with our results for Ase1, this raises the possibility that this short region within the carboxy-terminal domain provides dual function by binding importins and/or microtubules.

The distal region of the Ase1 carboxy-terminal domain (amino acids 694-885) recruits the EB1 homologue Bim1 to the midzone. Our results identify an SxIP motif at residues 775-778 of the carboxy-terminal domain of Ase1 that is required for this interaction. Sequence analysis suggests putative SxIP motifs in the carboxy-terminal domains of fission yeast Ase1 (SKSP, residues 614-617) and PRC1 (SRLP, residues 516-519; Supplemental Figure 1). Whether these motifs support interactions with EB1 homologues will require further investigation. Ablating the SxIP motif in budding yeast Ase1 (ase1^SHNN^) diminishes the recruitment of Bim1 to the midzone but does not impair Ase1 recruitment (Figure 5). These results add to the current model for Bim1 function in the anaphase spindle. Budding yeast mutants that lack Bim1 exhibit diminished anaphase spindle elongation forces, accompanied by significantly fewer interpolar microtubules (Gardner et al., 2008). This is consistent with Bim1 stabilizing and expanding the midzone to support anaphase spindle elongation. While Bim1 recruitment promotes anaphase spindle elongation, Bim1 inhibition is a key event in spindle disassembly at the end of anaphase. During late anaphase, Bim1 is phosphorylated by the budding yeast Aurora B kinase Ipl1 (Zimniak et al., 2009). This phosphorylation inhibits Bim1 microtubule-binding activity, removes it from the midzone, and promotes the timely disassembly of the spindle at the end of anaphase (Woodruff et al., 2010; Zimniak et al., 2009). These results suggest that Bim1’s recruitment to the midzone by Ase1, and subsequent eviction by Ipl1 phosphorylation are key determinants of anaphase spindle stability.

Given that Bim1 is expected to bind directly to the plus ends of interpolar microtubules, how might interaction with Ase1 contribute to its role in stabilizing the anaphase spindle? Our fluorescence intensity measurements suggest that interaction with Ase1 increases the amount of Bim1 at the midzone above what would be expected for plus-end binding alone and distributes Bim1 evenly across the midzone (Figure 5J and K). We propose that binding to Ase1 recruits Bim1 to microtubule lattices in the midzone, creating a ‘rescue zone’ where Bim1 could promote rescue of depolymerizing interpolar microtubules (Figure 6). This model is consistent with the established role of Bim1 in promoting rescue of astral microtubules during the G1 phase of the cell cycle (Tirnauer et al., 1999). By rescuing interpolar microtubules, Bim1 would maintain microtubule density and the extent of microtubule overlap in the midzone, and, therefore, increase the abundance of binding sites for Ase1. This would explain why depleting Bim1 from the midzone by ablating the SxIP motif in Ase1 leads to midzone compaction (Figure 5).

**Figure 6.**
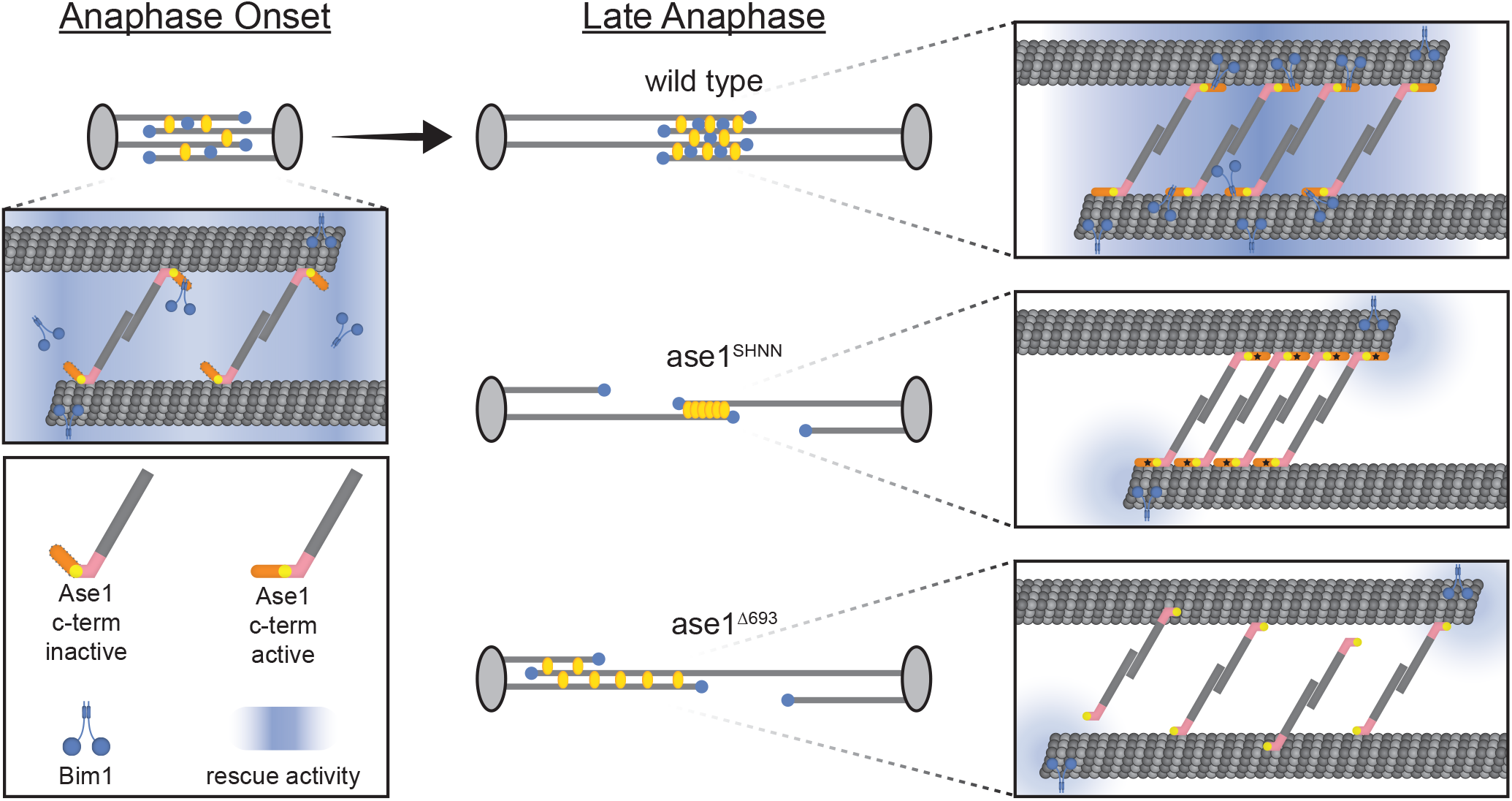
Model: Ase1 Domain Interactions Promote Modular Anaphase Function. Prior to and during anaphase onset Ase1 crosslinking is primarily driven by the spectrin domain. Bim1 is associated with the entire length of the short spindle. As the spindle progresses to late anaphase, the amount of Ase1 on the spindle increases and the carboxy-terminal tail contributes to slowing spindle elongation and maintaining Ase1’s association with microtubules. The Ase1 carboxy-terminal tail enriches the amount of Bim1 at the midzone and establishes a gradient of rescue activity throughout the midzone. In cells expressing *ase1^SHNN^* the interaction between the carboxy-terminal tail and Bim1 is ablated, and Bim1 only associates with the plus-ends of microtubules. This ablated ase1^SHNN^-Bim1 interaction results in fewer interpolar microtubules and greater compaction of ase1^SHNN^ within the midzone. In cells expressing *ase1*^*Δ693*^, Bim1 only associates with the plus-ends of microtubules and the processivity of ase1^Δ693^ on microtubules is decreased. This partial deletion of the carboxy-terminal domain results in an asymmetric midzone because of disrupted interactions between ase1^Δ693^ and microtubules or other proteins.

Bim1 binding is not the sole function of the distal region of Ase1’s carboxy-terminal domain. While ablating the Bim1-binding motif at residues 775-778 with the ase1^SHNN^ mutant causes midzone compaction, removing amino acids 694-885 from the carboxy-terminal domain (ase1^Δ693^) causes a more severe phenotype, including more frequent spindle collapses. Furthermore, the localization of ase1^SHNN^ is quite different from the ase1^Δ693^ truncation. Whereas ase1^SHNN^ localizes to a narrow region at the center of the anaphase spindle (Figure 5); ase1^Δ693^ localizes to a broader region and is not stably maintained at the center of the spindle (Figure 4). The corresponding carboxy-terminal domains of fission yeast Ase1, MAP65, and PRC1 exhibit direct microtubule-binding activity (Kapitein et al., 2008; Li et al., 2007; Mollinari et al., 2002; Smertenko et al., 2004; Subramanian et al., 2010). How these carboxy-terminal domains bind microtubules is still unclear since they are intrinsically disordered and refractory to structural studies; however, the data suggest that this most likely occurs through electrostatic interactions with microtubules (Kellogg et al., 2016; Subramanian et al., 2010). Electrostatic interactions of the carboxy-terminal domain could act as a processivity factor, in tandem with the spectrin domain, to keep Ase1 associated with midzone microtubules as they slide apart.

We propose a model in which Ase1 uses the distinct mechanisms of the spectrin and carboxy-terminal domains to support spindle elongation that is both fast and able to reach long lengths without collapse (Figure 6). Prior to anaphase, the spectrin domain promotes the recruitment of Ase1 to the spindle. During early anaphase, the spectrin domain binds and crosslinks microtubules, and spindle elongation reaches its fastest rate. The carboxy-terminal domain is dispensable during these phases. When the spindle reaches ~4.5 μm, approximately double its metaphase length, the lengths of interpolar microtubules become limiting and elongation begins to slow. From this point on, the carboxy-terminal domain of Ase1 sustains elongation through crosslinking interpolar microtubules and recruiting Bim1 to the midzone to prevent their depolymerization. Our model suggests a precisely timed activation of the carboxy-terminal domain in mid-anaphase, which could be achieved by phospho-regulation. Consistent with this notion, CDK1/Cdc28 phosphorylates Ase1 at 7 sites, including 4 CDK1 consensus sites in the carboxy-terminal domain (Khmelinskii et al., 2007, 2009; Ubersax et al., 2003) (Supplemental Figure 1). Cdc28 phosphorylates Ase1 during metaphase, inhibiting premature spindle elongation, while the Cdc14 phosphatase dephosphorylates Ase1 by mid-anaphase, which promotes spindle elongation (Khmelinskii et al., 2007, 2009). Interestingly, an *ase1* mutant that mimics constitutive phosphorylation at all 7 identified CDK sites exhibits asymmetric and broader distribution along the anaphase spindle, reminiscent of the ase1^Δ693^ truncation (Figure 4) (Khmelinskii et al., 2007). How phospho-regulation impacts the functions of the carboxy-terminal domain identified here, and how phospho-regulation might be coordinated with spindle length and midzone architecture will be an important area for future study.

## METHODS

### Yeast Strains and Manipulation

Chemicals and reagents were from Fisher Scientific (Pittsburgh, PA) and Sigma-Aldrich (Saint Louis, MO), unless stated otherwise. General yeast manipulation, media, and transformation were performed by standard methods (Amberg et al., 2005). Fluorescent tag fusions to Ase1, Bim1, Bik1 and Spc110 are integrated at the native loci (Sheff and Thorn, 2004). The mNeonGreen plasmid DNA was provided by Allele Biotechnology and Pharmaceuticals (San Diego, CA) (Shaner et al., 2013). GFP-Tub1 fusions were integrated at the *LEU2* locus and expressed ectopically to the native *TUB1* gene (Song and Lee, 2001). Mutant alleles of *ASE1* were generated using genomic DNA from the GFP tagged strain as a template. Regions containing the *ASE1* coding sequence, GFP, 3’UTR, and marker were amplified using mutagenic oligos, and transformed into a wild-type strain background. Mutations were confirmed by sequencing genomic loci, and were the only mutations present in the coding sequence. The *ase1*Δ deletion mutant was generated by conventional PCR-mediated methods (Petracek and Longtine, 2002). Strains and plasmids are listed in Supplementary Tables 3 and 4.

### Live Cell Imaging

Cells were grown overnight in rich media (2% glucose, 2% peptone, and 1% yeast extract) at 30°C then diluted into synthetic media (2% glucose, CSM from Sunrise Science Products, #1001 San Francisco, CA) and grown to log phase at 30°C. Cells were adhered to slide chambers coated with concanavalin A and sealed with VALAP (Vaseline, lanolin and paraffin at 1:1:1) (Fees et al., 2017). Images were collected on a Nikon Ti-E microscope equipped with a 1.45 NA 100× CFI Plan Apo objective, piezo electric stage (Physik Instrumente, Auburn, MA), spinning disk confocal scanner unit (CSU10; Yokogawa), 488-nm and 561-nm lasers (Agilent Technologies, Santa Clara, CA), and an EMCCD camera (iXon Ultra 897; Andor Technology, Belfast, UK) using NIS Elements software (Nikon).

During acquisition the temperature of the stage was 30°C for experiments (Figure 1, 2A-D, 4D-F, and 5) or 25°C (Figure 3 and 4B-C). Time lapse images were taken every 15 seconds (Figure 1 and 2) or every 10 seconds (Figure 4 and 5). Z series consisted of 13 images separated by 450 nm (Figure 1, 2A-D, 3, 4B-E, 5) or 11 images separated by 500 nm (Figure 4F).

### Spindle elongation analysis

Z-series images of cells expressing Spc110-tdTomato were first processed in FIJI (Schindelin et al., 2012) by using the “Despeckle” plugin followed by the “Remove Outlier” plugin with the radius set to 1 pixel and the threshold 25 to remove bright, outlier pixels. Spindle length over time was then determined using a custom MATLAB script that calculates the brightest pixel within the three-dimensional image stack, applies a gaussian blur around this pixel to subtract the effects of that SPB, and then identifies the second brightest pixel in the Z stack (i.e. the other SPB). Spindle length was defined as the linear distance between these two points in three dimensions. Any change in spindle length between frames that was greater than 500 nm was verified manually.

To identify elongation events, a custom MATLAB script was used to fit piecewise linear functions to the spindle length over time. First, the initial fast phase of anaphase was determined by identifying the largest change in spindle length within a 90 second period. Second, earlier time points were added one at a time to this region and a linear regression was calculated after each addition. The beginning of anaphase was defined as the earliest point that resulted in the highest r-squared value. From this starting point, a one-minute period (5 data points) was used to begin the piecewise process, and subsequent, single data points were added to this one-minute period and linear regressions were fit after each addition for up to 6.25 minutes (26 data points). The point resulting in the linear function with the last r-squared value greater than 0.95 was defined as the end of elongation event. If no linear regression resulted in a function with an r-squared value greater than 0.95, the addition of the data point resulting in the greatest r-squared value was defined as the end of the elongation event. The time point after the end was then used for a new one-minute period and the process repeated until all data was processed. The piecewise linear functions were then shown on top of the data points and verified by eye. Average r-squared values for all lines within their respective genotypes weighted by the number of data points included in the line: *ASE1−GFP* = 0.74, *ase1^3A^−GFP* = 0.75, *ase1^693Δ^−GFP* = 0.75, *ase1Δ* = 0.69.

The slope of the line from the piecewise linear regression process was then assigned as the elongation rate to all time points encompassed by that function. All values derived from linear regressions with r-squared values greater than 0.4 were then used for bins. The bins for time in anaphase were calculated equal to or greater than the minimum of the bin, and less than the maximum for one-minute increments (e.g., the bin for 1-2 minutes is graphed at bin center 1.5 min and is for 1 min ≤ values < 2 min). For the change in anaphase spindle length, the average spindle length of the two-minute period proceeding anaphase onset was subtracted from all spindle length values. Only the data for the first twenty minutes of anaphase were used for our analysis, in an attempt to exclude cells that may have exited mitosis. The bins were then calculated equal to or greater than the minimum of the bin, and less than the maximum for 500 nm increments.

### Analysis of spindle stability during telophase arrest

Cells were grown at room temperature to log phase and then shifted to 37°C. After 3 hours cells were fixed by incubation with fixative (0.5M KPO_4_ and 18.5% formaldehyde) for 2.5 minutes. Cells were then pelleted, washed with quench buffer (0.1M KPO_4_, 0.1% Triton-X, and 10mM ethanolamine), pelleted, and then washed with wash buffer (0.1M KPO_4_) and stored in wash buffer at 4°C until imaged. All cells were imaged within 24 hours of fixation. Images were collected on a Nikon Ti-E wide field microscope equipped with a 1.49 NA 100× CFI160 Apochromat objective, and an ORCA-Flash 4.0 LT sCMOS camera (Hammamatsu Photonics, Japan) using NIS Elements software (Nikon, Minato, Tokyo, Japan). Images were acquired in Z-series consisting of 13 images separated by 450 nm.

Cells were considered to have an intact spindle if there was a clear GFP signal connecting the two SPBs or aligned GFP signal that nearly connects the two SPBs. Otherwise, cells were considered to have a not intact spindle. Cells were only considered for analysis if there was observable fluorescence intensity above background and SPBs were separated by at least 2 μm.

### Ase1-GFP fluorescence intensity analysis

Z-series images of cells expressing Ase1-GFP and Spc110-tdTomato were sum projected and cytoplasmic background was subtracted, normalized to the area of the region of interest. The region of interest for experiments in Figure 3 and 5D was defined as a region extending between the two SPBs and including the Ase1-GFP signal with a width of 5 pixels (~300nm). Cytoplasmic background was determined by the average intensity of a sum-projected, 13-by-13-pixel box within the cell but adjacent to the spindle. Cells were only considered for analysis if the entire spindle was in focus as determined by the SPBs.

### Ase1-GFP midzone position analysis

Z-series images of cells expressing Ase1-GFP and Spc110-tdTomato were sum projected and cytoplasmic background was subtracted (as described above). The region of interest drawn for experiments in Figure 4 and 5E was defined a region extending from the SPB in the mother compartment to the SPB in the bud, with a width of 3 pixels. An average of the 3-pixel width along the region of interest was used to generate an intensity line scan, and the per pixel cytoplasmic background was subtracted from the line scan. A custom MATLAB script was used to determine the full-width half-max, and all results were verified by eye. The center of this Ase1 region was then identified, and its position along the region between the SPBs (from 0 at the mother SPB to 1.0 at the bud SPB) was determined.

### Bim1-tdTomato fluorescence intensity analysis

Images of cells expressing Ase1-GFP and Bim1-tdTomato were sum projected and cytoplasmic background was subtracted as described above. The region of interest drawn for experiments in Figure 5H-J was defined as a region between the SPBs estimated by Bim1-tdTomato signal and including the Ase1-GFP signal. An average of the 3-pixel width along the region of interest was used to generate an intensity line scan for both fluorescence channels, and the respective per-pixel cytoplasmic background was subtracted from the line scan for each channel. The length and midpoint of the Ase1 region was determined by the full-width half-max MATLAB script described above. The average Bim1-tdTomato intensity within the length of the Ase1-GFP region is reported in Figure 5H. To determine the fluorescence intensity of Bim1-tdTomato relative to the Ase1-GFP region an additional MATLAB script was used to identify two halves of each spindle, based the center of the Ase1 region. The intensities were recorded extending the length of the FWHM Ase1-GFP region, plus 50% of that length farther along the spindle and normalized to the total length, such that the edge of the Ase1 region was half-way along this line scan (see diagrams in Figure 5I and J).

To determine the ratio of total Bim1-tdTomato signal at the midzone relative to that on single astral microtubules (Figure 5K), a region of interest extending approximately 250nm on either side of the Ase1 region, with a 5-pixel width, was drawn by eye in anaphase cells. A 5-by-5-pixel box was drawn around foci of Bim1-tdTomato on clear astral microtubules within the same cell as the region of interest at the midzone. The cytoplasmic background was determined by the average intensity in a 13-by-13-pixel box within the cell adjacent to the spindle and any astral microtubules. Then the area-normalized background-subtracted Bim1 midzone intensity was divided by the area-normalized background-subtracted Bim1 astral microtubule foci intensity.

### Two-hybrid assay

Two-hybrid assay was conducted using Gal4-activation domain (pDEST-AD2) and Gal4-DNA-binding domain (pDEST-BD2) plasmids (a gift from Dr. Jay Hesselberth, University of Colorado School of Medicine). Full-length open reading frames were cloned from MORF plasmids (Gelperin et al., 2005) into pDEST-AD2 or pDEST-BD2 using Gateway cloning (ThermoFisher). For truncations of *ASE1*, regions were amplified from genomic DNA, cloned into a donor plasmid (pDONR221), and then into pDEST-BD2. Final fusion clones were confirmed by sequencing. To generate combinations of plasmids, pDEST-AD2 plasmids were transformed into yJM4404/PJ694α and pDEST-BD2 plasmids were transformed into yJM4403/PJ694A, and these haploid strains were mated to create diploids containing both plasmids. Diploid isolates were grown to late log phase in synthetic media lacking leucine and tryptophan (-Leu-Trp) to select for plasmid retention, spotted to media selective for plasmid retention (-Leu-Trp) or two-hybrid interaction (-Leu-Trp-His; -Leu-Trp-Ade) using a multi-prong transfer device, and grown at 30°C for 3 days before imaging. Empty pDEST-AD2 and pDEST-BD2 plasmids were used as controls.

### Statistics

Prism (GraphPad Software; San Diego, CA) was used for all graphs and statistical analysis, except for Supplemental Table 2, which used Microsoft Excel. For all multiple comparisons, an ANOVA test was first performed, and subsequent analysis was performed if p < 0.05. Student’s t-test was used for all homoscedastic, parametric data. Student’s t-test with Welch’s correction was used for heteroscedastic, parametric data. Mann-Whitney *U* test was used for nonparametric data. The Holm-Sidak t-test was used for the adjusted p-value in the multiple comparison data in Supplemental Table 1. The Microsoft Excel FTEST function was used for the F-test in Supplemental Table 2.

## ACKNOWLEDGEMENTS

We thank Dr. Elmar Schiebel for sharing plasmids for tagging Ase1 with GFP, Dr. Jay Hesselberth for sharing plasmids and yeast strains for the two-hybrid experiments, and members of the Moore lab for helpful discussions.

This work was supported by the National Institutes of Health grant R01 GM112893 to J.K. Moore. This material is based upon work supported by the National Science Foundation Graduate Research Fellowship under Grant No.1938058 to E.C. Thomas.

## ABBREVIATIONS

SPB: Spindle Pole Body
GFP: Green Fluorescent Protein
FWHM: Full-Width Half-Max
CI: Confidence Interval

## Notes

### Competing Interest Statement

The authors have declared no competing interest.

