## Supplemental Material for "Ase1 domains dynamically slow anaphase spindle elongation and recruit Bim1 to the midzone"

Fig. S1 shows a sequence alignment of Ase1 homologues across species.
Fig. S2 shows the localization of a panel if Ase1 truncations and point mutants to pre-anaphase spindles, and to the nucleus in cells treated with hydroxyurea and nocodazole.
Fig. S3 shows the full data set of spindle length changes measured in anaphase cells.
Fig. S4 shows cells expressing ase1∆693-GFP with signal on the spindle or without clear signal on the spindle.

Table S1 shows the full statistics for Figure 1.
Table S2 shows the full statistics for Figure 2.
Table S3 lists the strains used in this study.
Table S4 lists the plasmids used in this study.

Video 1 shows an example cell expressing Ase1-GFP and Spc110-tdTomato during early anaphase.
Video 2 shows an example cell expressing Ase1-GFP and Spc110-tdTomato during late anaphase.
Video 3 shows an example cell expressing ase1$^{\Delta 693}$-GFP and Spc110-tdTomato during early anaphase.
Video 4 shows an example cell expressing ase1$^{\Delta 693}$-GFP and Spc110-tdTomato during late anaphase.

##### Supplemental Figure 1 – Sequence Alignment

Alignment of S. cerevisiae Ase1 (CAA99251.1), S. pombe Ase1 (NP_593523.1), H. sapiens PRC1 (AAC02688.1), and A. thalania MAP65-1 (NP_001190542.1) generated by Clustal Omega. Gray region denotes spectrin domain (aa523-650 of Ase1). Yellow region denotes proximal carboxy-terminal domain (aa651-693 of Ase1). Orange region denotes distal carboxy-terminal domain (aa694-885). Residues highlighted in green are identified CDK phosphorylation sites (Khmelinski1, et al. 2007). Blue residues are predicted to interact with the β-tubulin carboxy-terminal tail (Kellogg, et al. 2016). Pink region denotes the Bim1-binding motif (aa775-778).

##### Supplemental Figure 2 – Expression and pre-anaphase localization of ase1 mutants

A) Domain architecture of PRC1 and Ase1 with ase1 mutant constructs.
B) Example images of wild-type Ase1-GFP in pre-anaphase. Scale bar = 1 µm.
C) Cells were arrested in S-phase with a short bipolar spindle by treatment with 220mM hydroxyurea in YPD for two hours. Cells were then treated with an additional 20uM nocodazole for an hour to depolymerize spindle microtubules. Fluorescence analysis was performed as described in Materials and Methods with a region of interest drawn around the nucleus as determined by overexposure of the Spc110-tdTomato signal. Ase1-GFP, *n* = 60; No GFP, *n* = 63; ase1$^{SC}$-GFP, *n* = 60; ase1$^{\Delta 650}$, *n* = 54; ase1$^{\Delta 693}$, *n* = 55; ase1$^{3A}$, *n* = 44.
D) Intensity of Ase1-GFP on pre-anaphase spindles was measured in asynchronous cells with a spindle length ≤ 1.5 µm. 2-dimensional projections of Z-series images were created by sum projection, and total GFP intensity values were measured in an ROI covering the area between the SPBs.
E) Pre-anaphase spindle lengths were determined from the same images as in (D).

F) Ratio of pre-anaphase spindle intensity to length was determined by dividing the total Ase1-GFP signal intensity (D) by the length of the spindle in the same cell (E).
G) Example images of hydroxyurea nocodazole treated cells used for the analysis in C. Scale bar = 1 μm.
*p<0.05, **p<0.01, determined by Student's t-test.

**Supplemental Figure 3 – Full data set for anaphase spindle elongation**

A) Average anaphase rates binned by one-minute increments from anaphase onset. Data from wild-type controls is shown in black in each graph for comparison. Error bars represent 95% CI of the mean.
B) Average anaphase rates binned by half-micron increments of change in spindle length from anaphase onset. Data from wild-type controls is shown in black in each graph for comparison. Error bars represent 95% CI of the mean.
C) The average spindle length of the two minutes preceding anaphase onset. Bars indicate median ± 95% CI.
D) The average spindle length versus time for each genotype. Dotted lines represent linear regressions fit to the first three minutes or minutes 5-10 since anaphase onset. The transition from the fast to slow phase of anaphase was defined as the spindle length of the intersection of the two linear regression models. Ase1, $n$ = 10; ase1$^{3A}$, $n$ = 15; ase1$^{\Delta 693}$, $n$ = 11; ase1Δ, $n$ = 14.

**Supplemental Figure 4 – Quantification of diffuse nuclear Ase1 signal**

A) Example images of cells that appear to be in anaphase based on the positioning of the SPBs and have increased GFP fluorescence over than of the cytoplasmic background. Scale bar = 1 μm.
B) Percentage of cells that appear to be in anaphase but have diffuse nuclear GFP fluorescence. Error bars represent mean ± 95% CI.

**Supplemental Table 1 – Full statistics for Figure 1 panels H and I showing elongation rates**

P-value shown from Student's t-test to wild-type control. Adjusted p-value from Holm-Sidak test for multiple comparisons to wild-type control.

**Supplemental Table 2 – Full statistics for Figure 2 panels B and C showing average spindle length over time and standard deviation**

F-test for variance reports two-tailed probability for comparison to wild-type control.

**Supplemental Table 3 – Strains used in this study**

**Supplemental Table 4 – Plasmids used in this study**

**Supplemental Video 1 – Example wild-type cell from Figure 4D as it progresses through early anaphase.** Spc110-tdTomato (magenta) and Ase1-GFP (green). Frame rate = 10 frames per second. Scale bar = 1 μm.

**Supplemental Video 2 – Example wild-type cell from Figure 4D as it progresses through mid-to-late anaphase.** Spc110-tdTomato (magenta) and Ase1-GFP (green). Frame rate = 10 frames per second. Scale bar = 1 μm.

**Supplemental Video 3 – Example *ase1$^{\Delta 693}$* cell from Figure 4D as it progresses through early anaphase.** Spc110-tdTomato (magenta) and ase1$^{\Delta 693}$-GFP (green). Frame rate = 10 frames per second. Scale bar = 1 μm.

**Supplemental Video 4 – Example *ase1$^{\Delta 693}$* cell from Figure 4D as it progresses through mid-to-late anaphase.** Spc110-tdTomato (magenta) and ase1$^{\Delta 693}$-GFP (green). Frame rate = 10 frames per second. Scale bar = 1 μm.

### Supplemental Figure 1.

- spectrin domain
- proximal carboxy-terminal domain
- distal carboxy-terminal domain
- CDK phosphorylation sites
- basic residues mutated in ase13A
- SHIP motif

```
Sc_Ase1      METATSSPLPIKSRRNSENSGSTTVIPHMNPSLATPLTVSTMVNQSNSKEFMKLTPVRIR 60
Sp_ase1      -----------------------------------------------------MQTVMMD 7
Hs_PRC1      ------------------------------------------------------------ 0
At_MAP65-1   ------------------------------------------------------------ 0

Sc_Ase1      DFGSPLKNVSTNYHFLDSENGKGNTMDNMYRENFILISKDLEKLLENLNVIYQNIGYSNT 120
Sp_ase1      DIQS-----------TDSIAEKDNHSNNESNFTWKAFREQVEKHFSKIERLHQVLGTDGD 56
Hs_PRC1      --------------------MRRSEVLAEESIV------CLQKALNHLREIWELIGIPED 34
At_MAP65-1   ------------------MAVTDTESPHLGEIT-------CGTLLEKLQEIWDEVGESDD 35
                                    .                  . :.::. : : :*

Sc_Ase1      EIITKEKIIFTTISNSIKQF--------------FEQADEELKRLSAENGIEQDILNNIL 166
Sp_ase1      NSSL---------FELFTTA-------MNAQLHEMEQCQKKLEDDCQQRIDSIRFLVSSL 100
Hs_PRC1      QRLQRTEVVKKHIKELLDMMIAEEESLKERLIKSISVCQKELNTLCSELHVE------P- 87
At_MAP65-1   ERDKLLLQIEQECLDVYKRKVEQAAKSRAELLQTLSDANAELSSLTMSLGDK------SL 89
             :             :                   :. .: :*.    .   .

Sc_Ase1      ERINDPSGIKTIPDLYIRNAILLQESKTVPQSPKKPLSLLSKKAALDTAKKFVLGSFLPR 226
Sp_ase1      KLTDDTSSLK------IESPLI------------------QCLNRL----SMVEGQYMAQ 132
Hs_PRC1      --FQEEGE----------TTIL------------------QLEKDL----RTQVELMRKQ 113
At_MAP65-1   VGIPDKSS----------GTIK------------------EQLAAI----APALEQLWQQ 117
                : .             :                   .    :             :

Sc_Ase1      LRDYLKSLITLKHLIQSVKENLPGLTEADNEAIAEFPELSTLTAYLLQIENGKGDIGLSM 286
Sp_ase1      YDQKLSTIKEMYHKLESYCNRLGSPFVLP------------------------------- 161
Hs_PRC1      KKERKQELKLLQEQDQELCEILCMPH---------------------------------- 139
At_MAP65-1   KEERVREFSDVQSQIQKICGDIAGGLSNE------------------------------- 146
               :    :  :    :.    :

Sc_Ase1      KFIIDNRKDILKGSAFKTINEESVKHMNEVIKIYEEEYERRFKSVLTKKVSISSICEQLG 346
Sp_ase1      DFENSFLSD---------VSDAFTESLRGRINEAEKEIDARLEVINSFEEEILGLWSELG 212
Hs_PRC1      -YDIDSASV---------PSLEELNQFRQHVTTLRETKASRRE---EFVSIKRQIIL--C 184
At_MAP65-1   -VPIVDESD---------LSLKKLDDFQSQLQELQKEKSDRLRKVLEFVSTVHDLCAVLG 196
                    .           .    . :.  :    .:    * .           :

Sc_Ase1      TPLATLIGEDFEQDLRSYGEEE-NSTSEIPNFHPVDRERMSKIDITLEKLQAIHKERADK 405
Sp_ase1      VEPADVP--QYEQLLESHTNRPN--------DVYVTQELIDQLCKQKEVFSAEKEKRSDH 262
Hs_PRC1      MEELDH-------T--PDTSFERDVVCEDEDAFCLSLENIATLQKLLRQLEMQKSQNEAV 235
At_MAP65-1   LDFLST-------VTEVHPSLDEDTSVQ---SKSISNETLSRLAKTVLTLKDDKKQRLQK 246
                           .              :  * :  :      :.  :.:.

Sc_Ase1      KRLLMEQCQKLWTRLKISQEYIKTFMRNNSSLST-------------------------- 439
Sp_ase1      LKSIQSEVSNLWNKLQVSPNEQSQFGDSSN-----INQ---------------------- 295
Hs_PRC1      CEGLRTQIRELWDRLQIPEEEREAVATIMSGSKAKVRK---------------------- 273
At_MAP65-1   LQELATQLIDLWNLMDTPDEERELFDHVTCNISSSVDEVTVPGALARDLIEQVIYIALIN 306
              . :  :  .**  :.    :  .  .

Sc_Ase1      ----------------ESLGRISKEVMRLEAMKKKLIKKLISDSWDKIQELWRTLQYSEE 483
Sp_ase1      ----------------ENISLWETELEKLHQLKKEHLPIFLEDCRQQILQLWDSLFYSEE 339
Hs_PRC1      --------------------ALQLEVDRLEELKMQNMKKVIEAIRVELVQYWDQCFYSQE 313
At_MAP65-1   LPMSSLRNQLLLANIHVKFVKAEVEVDRLDQLKASRMKEIAFKKQSELEEIYARAHVEVN 366
                                     . *: :*. :* . :   .      :: : :    . :

Sc_Ase1      ---SRSKFIIVFEELRNSATTLQEDELLLETCENELKRLEEKLTLYKPILKLISDFESLQ 540
Sp_ase1      ---QRKSFTPMYEDII--------TEQVLTAHENYIKQLEAEVSANKSFLSLINRYASLI 388
Hs_PRC1      ---QRQAFAPFCAEDY--------TESLLQLHDAEIVRLKNYYEVHKELFEGVQKWEETW 362
At_MAP65-1   PESARERIMSLIDSGN------VEPTELLADMDSQISKAKEEAFSRKDILDRVEKWMSAC 420
                *. :   .   .             :*   :   : : :      * ::. :. : .

Sc_Ase1      EDQEFLERSSKDSSRLLSRNS--HKILLTEEKMRKRITRHFPKVINDLRIKLEEADGLFD 598
Sp_ase1      EGKKELEASSNDASRLTQRGRRDPGLLLREEKIRKRLSRELPKVQSLLIPEITAWEERNG 448
Hs_PRC1      RLFLEFERKASDPNRFTNRGG--NLLK--EEKQRAKLQKMLPKLEEELKARIELWEQEHS 418
At_MAP65-1   EEESWLEDYNRDQNRYSASRG--AHLNLKRAEKARILVSKIPAMVDTLVAKTRAWEEEHS 478
             .    :*     *  .*             :    . :     :    :* : . *    .     :    .

Sc_Ase1      QPFLFKGKPLSEAIDIQQQEIEAKYPR----CRVRMQRSKKG-KCGANKENKVIKNTFKA 653
Sp_ase1      RTFLFYDEPLLKICQEATQPKSLYRSASAAANRPKTATTDSVNRTPSQRGRV---AVPS 505
Hs_PRC1      KAFMVNGQKFMEYVAEQWEMHRLEKERAKQERQLKN---KK------QTETEM---LYGS 466
At_MAP65-1   MSFAYDGVPLLAMLDE-YGMLRQEREE--EKRRLRE---QKKVQEQPHVEQES---AFST 529
               *    .   :                    : :       .          . .        :

Sc_Ase1      TESSIRVPIGLNLNDANITYKTPSKKTIQGLTKNDLSQENSLARHMQGTTKLSSPNRRAT 713
Sp_ase1      TPSVRSASRAMT------SPRTPLP-RVKNT---------------------QNPSR--- 534
Hs_PRC1      APRTPSKRRGLA-------PNTPGKARKLNTTTMSNATANSSIRPIFGGTVYHSPVSRLP 519
At_MAP65-1   RPSP-------------------------------------------------------- 533

Sc_Ase1      RLLAPTVISRNSKGNIERPT------------------------LNRNRS--SDLSSSP 746
Sp_ase1      SISAEPPSATSTAN-RRHPTANRIDINARLNSASRSRSANMIRQGANGSDS---NMSSSP 590
Hs_PRC1      PSGSKPVAASTCSG-KKTP----------------------RTGRHGANKENLELNGSI 555
At_MAP65-1   ---ARPVSAKKTVG-PRAN----------------------NGGANGTHNRRLSLNANQ 566
                :    : .  .   .                                    :    .    .:...

Sc_Ase1      RINHTHGEHAVKPRQLFPIPLNKVDTKGSH-----IPQLTKEK----------------- 784
Sp_ase1      VSG-----NSNTPFNKFPNSVSRNTHFESKSPHPNYSRTPHETYS------KASSKNVPL 639
Hs_PRC1      LSG-----GYP-----GSAPLQRN--FSINSVASTYSEFAKDPSLSDSSTVGLQRELSKA 603
At_MAP65-1   NGS-----RST------AKEAGRRETLNRPAAPTNYVAISKEEAASSP-VSGAADHQVPA 614
               .                   :                 ::

Sc_Ase1      -----------ALELLKRSTGTTGKENVRSPERKSSLEDYAQKLS------SPYKE-PEH 826
Sp_ase1      SPPKQRVVNEHALNIM---SEKLQRTNLKEQTPEMDIENSSQNLPFSPMKISPIRASPVK 696
Hs_PRC1      SKSDAT-------------SGILNSTNIQS------------------------------ 620
At_MAP65-1   SP---------------------------------------------------------- 616

Sc_Ase1      SIYKLSMSPEGKFQLNIQQKDIESGFDDTSMMEDENDKDFITWKNEQVSKLNGFSFTDI 885
Sp_ase1      T---IPSSPSPT--TNIFSAPLN-NITNCTPMED-------EWGEEGF----------- 731
Hs_PRC1      ----------------------------------------------------------- 620
At_MAP65-1   ----------------------------------------------------------- 616
```

### Supplemental Figure 2 - Expression and pre-anaphase localization of *ase1* mutants

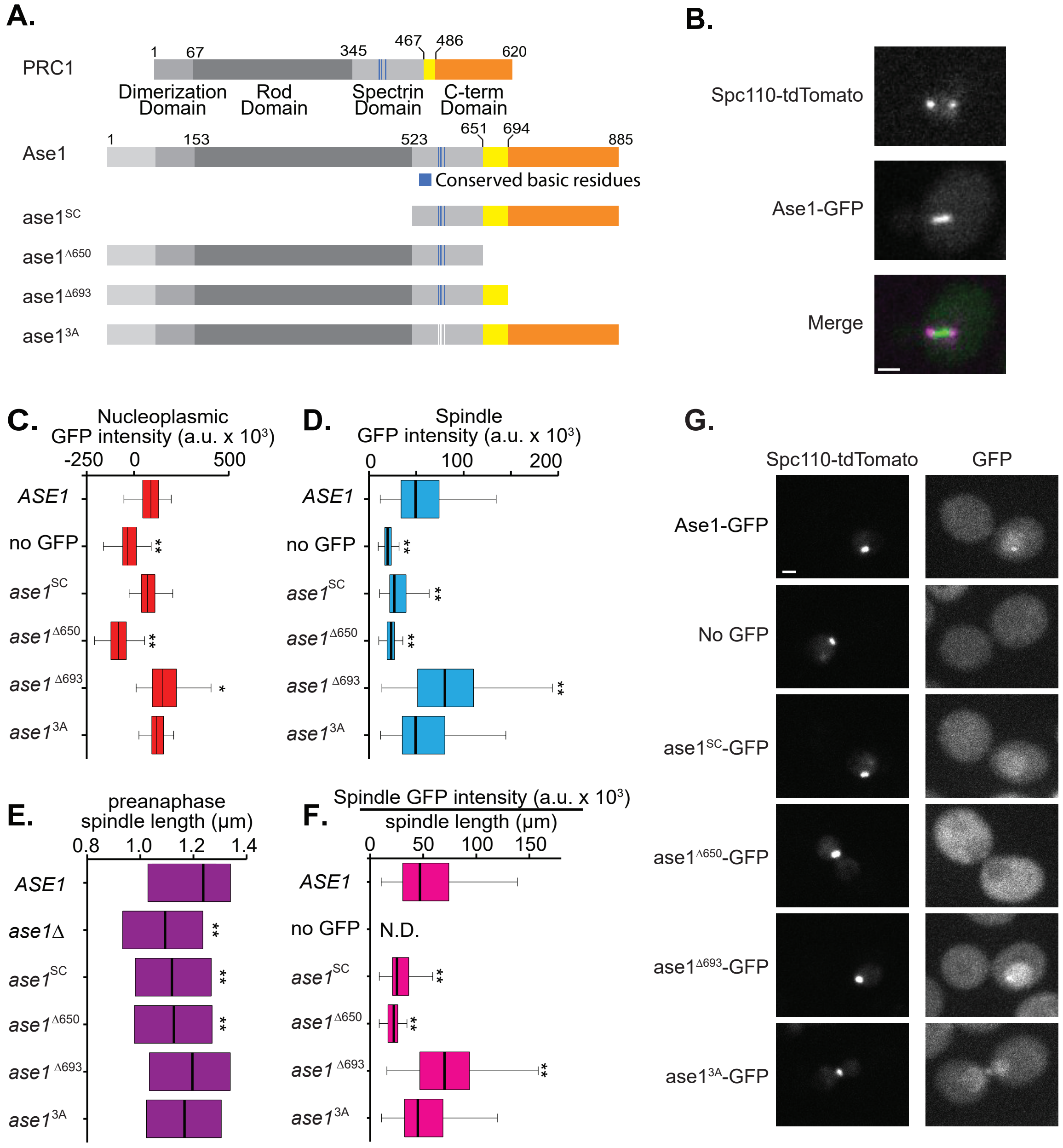

### Supplemental Figure 3 - Full data set for anaphase spindle elongation

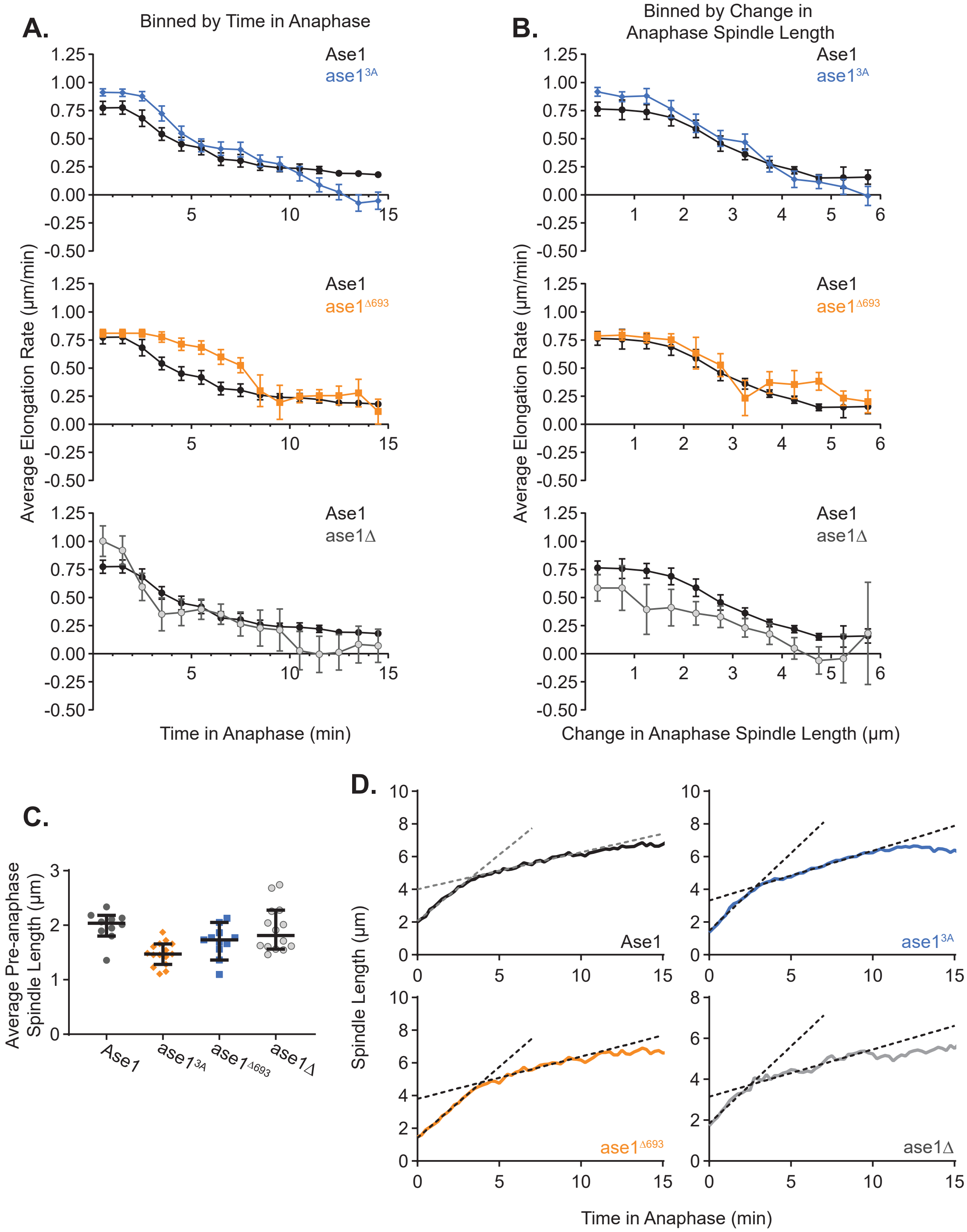

#### Supplemental Figure 4 - Quantification of diffuse nuclear Ase1 signal

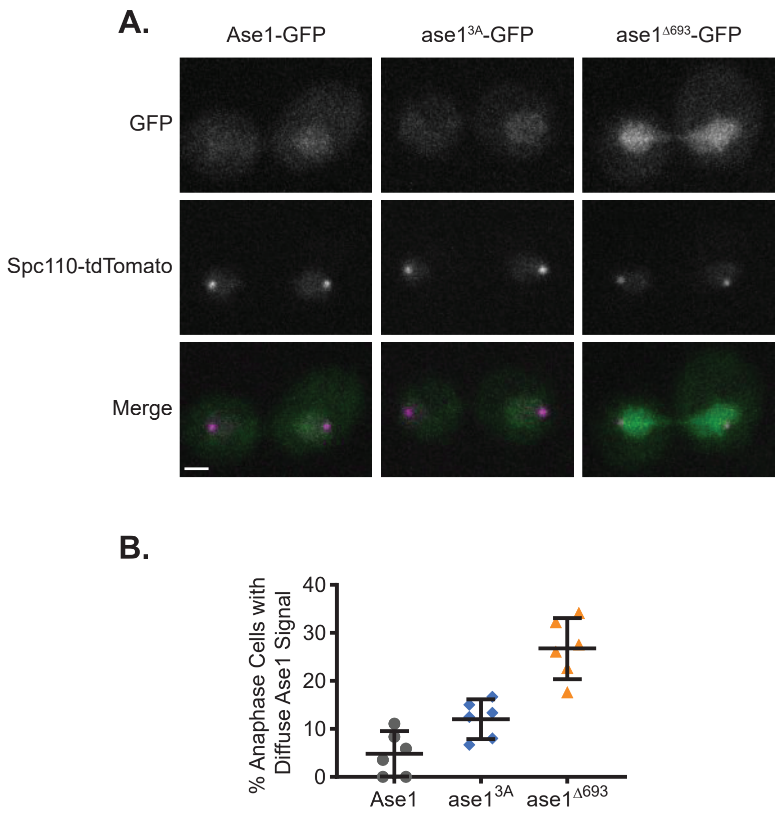

| Binned by Time in Anaphase | | | | | | | | | | | | | | | | | |
|---|---|---|---|---|---|---|---|---|---|---|---|---|---|---|---|---|---|
| | Ase1 | | | ase1$^{3A}$ | | | | | ase1$^{\Delta 693}$ | | | | | ase1Δ | | | |
| Bin Center (min) | avg | sd | n | avg | sd | n | p | adjusted p | avg | sd | n | p | adjusted p | avg | sd | n | p |
| 0.5 | 0.774 | 0.187 | 39 | 0.912 | 0.129 | 60 | 4E-05 | 0.0005 | 0.81 | 0.116 | 44 | 0.298 | 0.5071 | 1.002 | 0.519 | 56 | 0.01 |
| 1.5 | 0.776 | 0.186 | 39 | 0.909 | 0.128 | 59 | 6E-05 | 0.0008 | 0.81 | 0.116 | 44 | 0.317 | 0.5071 | 0.92 | 0.481 | 55 | 0.079 |
| 2.5 | 0.681 | 0.233 | 40 | 0.878 | 0.164 | 60 | 3E-06 | 5E-05 | 0.81 | 0.116 | 44 | 0.002 | 0.0071 | 0.674 | 0.308 | 50 | 0.902 |
| 3.5 | 0.54 | 0.181 | 40 | 0.724 | 0.256 | 58 | 2E-04 | 0.0021 | 0.776 | 0.154 | 43 | <0.00000 | <0.000001 | 0.561 | 0.173 | 46 | 0.583 |
| 4.5 | 0.452 | 0.192 | 39 | 0.549 | 0.233 | 55 | 0.035 | 0.2678 | 0.713 | 0.18 | 42 | <0.00000 | <0.000001 | 0.504 | 0.18 | 48 | 0.2 |
| 5.5 | 0.418 | 0.198 | 40 | 0.441 | 0.222 | 58 | 0.599 | 0.9356 | 0.684 | 0.198 | 43 | <0.00000 | <0.000001 | 0.473 | 0.198 | 44 | 0.205 |
| 6.5 | 0.318 | 0.181 | 40 | 0.411 | 0.231 | 58 | 0.034 | 0.2678 | 0.6 | 0.205 | 38 | <0.00000 | <0.000001 | 0.439 | 0.197 | 45 | 0.004 |
| 7.5 | 0.304 | 0.173 | 38 | 0.403 | 0.251 | 55 | 0.038 | 0.2678 | 0.524 | 0.226 | 44 | 5E-06 | 5E-05 | 0.414 | 0.181 | 44 | 0.006 |
| 8.5 | 0.259 | 0.128 | 39 | 0.33 | 0.141 | 54 | 0.015 | 0.1515 | 0.44 | 0.235 | 37 | 7E-05 | 0.0006 | 0.472 | 0.33 | 37 | 4E-04 |
| 9.5 | 0.24 | 0.1 | 37 | 0.315 | 0.176 | 50 | 0.023 | 0.2044 | 0.374 | 0.187 | 35 | 3E-04 | 0.0018 | 0.46 | 0.382 | 31 | 0.001 |
| 10.5 | 0.236 | 0.113 | 36 | 0.275 | 0.144 | 42 | 0.194 | 0.66 | 0.304 | 0.082 | 29 | 0.009 | 0.028 | 0.307 | 0.116 | 23 | 0.024 |
| 11.5 | 0.221 | 0.099 | 36 | 0.223 | 0.09 | 34 | 0.92 | 0.9796 | 0.307 | 0.081 | 32 | 2E-04 | 0.0017 | 0.315 | 0.126 | 19 | 0.004 |
| 12.5 | 0.191 | 0.059 | 33 | 0.212 | 0.081 | 27 | 0.252 | 0.6874 | 0.332 | 0.175 | 28 | 5E-05 | 0.0005 | 0.413 | 0.157 | 17 | <0.00000 |
| 13.5 | 0.189 | 0.059 | 30 | 0.185 | 0.077 | 18 | 0.857 | 0.9796 | 0.424 | 0.285 | 28 | 5E-05 | 0.0004 | 0.386 | 0.167 | 23 | <0.00000 |
| 14.5 | 0.179 | 0.063 | 28 | 0.231 | 0.148 | 21 | 0.103 | 0.4792 | 0.284 | 0.132 | 21 | 6E-04 | 0.003 | 0.313 | 0.128 | 18 | 2E-05 |

| Binned by Change in Anaphase Spindle Length | | | | | | | | | | | | | | | | | |
|---|---|---|---|---|---|---|---|---|---|---|---|---|---|---|---|---|---|
| | Ase1 | | | ase1$^{3A}$ | | | | | ase1$^{\Delta 693}$ | | | | | ase1Δ | | | |
| Bin Center (μm) | avg | sd | n | avg | sd | n | p | adjusted p | avg | sd | n | p | adjusted p | avg | sd | n | p |
| 0.25 | 0.764 | 0.168 | 30 | 0.917 | 0.128 | 44 | 3E-05 | 0.0003 | 0.785 | 0.112 | 48 | 0.494 | 0.7958 | 0.718 | 0.388 | 80 | 0.537 |
| 0.75 | 0.756 | 0.177 | 16 | 0.874 | 0.117 | 27 | 0.012 | 0.0604 | 0.794 | 0.12 | 27 | 0.411 | 0.7958 | 0.822 | 0.396 | 38 | 0.53 |
| 1.25 | 0.738 | 0.167 | 25 | 0.879 | 0.162 | 23 | 0.005 | 0.0271 | 0.77 | 0.114 | 26 | 0.419 | 0.7958 | 0.73 | 0.322 | 37 | 0.916 |
| 1.75 | 0.688 | 0.197 | 27 | 0.763 | 0.257 | 44 | 0.2 | 0.4889 | 0.752 | 0.134 | 24 | 0.186 | 0.5612 | 0.646 | 0.387 | 45 | 0.597 |
| 2.25 | 0.586 | 0.231 | 34 | 0.637 | 0.346 | 70 | 0.443 | 0.5874 | 0.731 | 0.164 | 42 | 0.002 | 0.0128 | 0.478 | 0.351 | 81 | 0.102 |
| 2.75 | 0.456 | 0.269 | 61 | 0.503 | 0.317 | 80 | 0.358 | 0.5874 | 0.621 | 0.244 | 60 | 6E-04 | 0.0045 | 0.491 | 0.378 | 110 | 0.524 |
| 3.25 | 0.36 | 0.204 | 74 | 0.495 | 0.284 | 74 | 0.001 | 0.0081 | 0.518 | 0.277 | 40 | 7E-04 | 0.0052 | 0.404 | 0.291 | 106 | 0.267 |
| 3.75 | 0.286 | 0.147 | 99 | 0.419 | 0.257 | 99 | 1E-05 | 0.0001 | 0.554 | 0.26 | 65 | <0.00000 | <0.000001 | 0.392 | 0.299 | 81 | 0.002 |
| 4.25 | 0.23 | 0.154 | 108 | 0.39 | 0.194 | 80 | <0.00000 | <0.000001 | 0.493 | 0.279 | 49 | <0.00000 | <0.000001 | 0.28 | 0.159 | 44 | 0.07 |
| 4.75 | 0.181 | 0.132 | 127 | 0.343 | 0.225 | 101 | <0.00000 | <0.000001 | 0.449 | 0.258 | 57 | <0.00000 | <0.000001 | 0.333 | 0.211 | 23 | 1E-05 |

| | | | | | | | | | | | | | | | | | |
|---|---|---|---|---|---|---|---|---|---|---|---|---|---|---|---|---|---|
| 5.25 | 0.211 | 0.137 | 31 | 0.323 | 0.234 | 56 | 0.017 | 0.065 | 0.286 | 0.232 | 62 | 0.1 | 0.4089 | 0.44 | 0.164 | 14 | 1E-05 |
| 5.75 | 0.211 | 0.081 | 33 | 0.382 | 0.258 | 32 | 6E-04 | 0.0046 | 0.382 | 0.211 | 37 | 4E-05 | 0.0004 | 0.59 | 0.061 | 3 | <0.00000 |

| adjusted p |
|---|
| 0.0684 |
| 0.3365 |
| 0.9022 |
| 0.8261 |
| 0.5901 |
| 0.5901 |
| 0.036 |
| 0.0484 |
| 0.0044 |
| 0.0136 |
| 0.1359 |
| 0.0352 |
| <0.000001 |
| 3E-06 |
| 0.0003 |

| adjusted p |
|---|
| 0.9757 |
| 0.9757 |
| 0.9757 |
| 0.9757 |
| 0.5285 |
| 0.9757 |
| 0.8446 |
| 0.0204 |
| 0.4383 |
| 0.0001 |

| |
|---|
| 0.0001 |
| <0.000001 |

| | Ase1 | | | $ase1^{3A}$ | | | | $ase1^{\Delta693}$ | | | | ase1Δ | | | |
|---|---|---|---|---|---|---|---|---|---|---|---|---|---|---|---|
| Time (min) | Avg | SD | n | Avg | SD | n | F-test | Avg | SD | n | F-test | Avg | SD | n | F-test |
| 0 | 2.0122 | 0.2199 | 10 | 1.3732 | 0.2918 | 15 | 0.3965 | 1.4724 | 0.21 | 11 | 0.8814 | 1.728 | 0.4491 | 14 | 0.038 |
| 0.25 | 2.2012 | 0.3092 | 10 | 1.6293 | 0.3722 | 15 | 0.5849 | 1.5911 | 0.297 | 11 | 0.8941 | 1.9574 | 0.5256 | 14 | 0.1165 |
| 0.5 | 2.3379 | 0.2957 | 9 | 1.8639 | 0.3088 | 15 | 0.9388 | 1.9062 | 0.2361 | 11 | 0.4958 | 2.138 | 0.7043 | 14 | 0.0193 |
| 0.75 | 2.7724 | 0.4242 | 10 | 2.174 | 0.4273 | 15 | 0.9836 | 2.0797 | 0.4879 | 11 | 0.6844 | 2.438 | 0.7467 | 14 | 0.0953 |
| 1 | 2.9276 | 0.4 | 10 | 2.4166 | 0.4945 | 15 | 0.5285 | 2.2397 | 0.6397 | 11 | 0.1732 | 2.6609 | 0.9619 | 13 | 0.013 |
| 1.25 | 3.1607 | 0.4096 | 10 | 2.8792 | 0.4594 | 14 | 0.7451 | 2.4221 | 0.5526 | 11 | 0.3813 | 2.947 | 0.9533 | 14 | 0.0157 |
| 1.5 | 3.4105 | 0.4573 | 9 | 3.1236 | 0.4247 | 15 | 0.7722 | 2.7523 | 0.6559 | 11 | 0.3189 | 3.1243 | 1.0583 | 14 | 0.0231 |
| 1.75 | 3.5542 | 0.5123 | 10 | 3.2265 | 0.4189 | 15 | 0.4818 | 2.9043 | 0.725 | 11 | 0.3108 | 3.2912 | 1.082 | 14 | 0.0305 |
| 2 | 3.6985 | 0.488 | 10 | 3.4087 | 0.3908 | 15 | 0.4397 | 3.1331 | 0.8047 | 11 | 0.1479 | 3.3283 | 1.1576 | 14 | 0.0138 |
| 2.25 | 3.8824 | 0.4872 | 10 | 3.5939 | 0.3643 | 15 | 0.3179 | 3.4475 | 0.9105 | 11 | 0.0733 | 3.4604 | 1.0168 | 14 | 0.033 |
| 2.5 | 4.1503 | 0.512 | 10 | 3.8489 | 0.4164 | 15 | 0.4706 | 3.5406 | 0.7686 | 11 | 0.237 | 3.4715 | 0.9053 | 12 | 0.0978 |
| 2.75 | 4.2027 | 0.465 | 10 | 3.9653 | 0.4546 | 15 | 0.9068 | 3.8303 | 0.8524 | 11 | 0.0821 | 4.078 | 1.1002 | 13 | 0.0146 |
| 3 | 4.4018 | 0.4068 | 10 | 4.1913 | 0.5491 | 15 | 0.3682 | 4.0326 | 0.8384 | 11 | 0.0403 | 4.008 | 1.1024 | 14 | 0.0052 |
| 3.25 | 4.5682 | 0.3782 | 10 | 4.3422 | 0.5798 | 15 | 0.2002 | 4.2447 | 0.8189 | 11 | 0.0293 | 3.9593 | 1.1258 | 13 | 0.0027 |
| 3.5 | 4.6256 | 0.3597 | 10 | 4.4482 | 0.6751 | 15 | 0.0633 | 4.4406 | 0.8233 | 10 | 0.0215 | 4.2333 | 1.0685 | 14 | 0.0026 |
| 3.75 | 4.8011 | 0.3325 | 10 | 4.5056 | 0.6773 | 15 | 0.0376 | 4.5559 | 0.95 | 11 | 0.0041 | 3.9756 | 1.0369 | 13 | 0.0019 |
| 4 | 4.801 | 0.3197 | 10 | 4.6619 | 0.6813 | 15 | 0.0278 | 4.6662 | 0.7959 | 9 | 0.0131 | 4.1775 | 1.0593 | 14 | 0.0011 |
| 4.25 | 4.9143 | 0.3964 | 10 | 4.6314 | 0.6784 | 15 | 0.1099 | 4.748 | 0.8207 | 11 | 0.0391 | 4.1754 | 1.2256 | 14 | 0.0019 |
| 4.5 | 4.9444 | 0.3113 | 10 | 4.6013 | 0.6464 | 14 | 0.0341 | 4.8976 | 1.0353 | 11 | 0.0013 | 4.3005 | 1.3577 | 14 | 0.0001 |
| 4.75 | 5.0996 | 0.2621 | 9 | 4.7345 | 0.7063 | 15 | 0.0083 | 4.8693 | 0.97 | 11 | 0.0011 | 4.3539 | 1.2208 | 14 | 0.0002 |
| 5 | 5.0307 | 0.2683 | 10 | 4.7622 | 0.7521 | 15 | 0.0039 | 4.7608 | 1.0178 | 10 | 0.0005 | 4.4688 | 1.1902 | 14 | 0.0001 |
| 5.25 | 5.2718 | 0.3518 | 10 | 4.894 | 0.7898 | 15 | 0.0195 | 5.0199 | 1.094 | 11 | 0.0022 | 4.4459 | 1.0232 | 12 | 0.0034 |
| 5.5 | 5.203 | 0.3299 | 10 | 4.9916 | 0.8561 | 15 | 0.0069 | 5.3139 | 1.0558 | 11 | 0.0018 | 4.3482 | 0.8775 | 12 | 0.0066 |
| 5.75 | 5.238 | 0.3534 | 10 | 5.0099 | 0.824 | 15 | 0.015 | 5.246 | 1.1038 | 11 | 0.0021 | 4.343 | 0.8131 | 12 | 0.0185 |
| 6 | 5.3095 | 0.5737 | 10 | 5.1573 | 0.851 | 15 | 0.2365 | 5.4311 | 1.1041 | 10 | 0.0643 | 4.4062 | 0.867 | 13 | 0.2215 |
| 6.25 | 5.4462 | 0.397 | 10 | 5.2849 | 0.8339 | 15 | 0.0307 | 5.5002 | 1.068 | 9 | 0.0076 | 4.3516 | 0.8124 | 13 | 0.0388 |
| 6.5 | 5.6231 | 0.4404 | 10 | 5.2977 | 0.847 | 14 | 0.0558 | 5.7467 | 1.0213 | 9 | 0.0211 | 4.5328 | 1.0562 | 14 | 0.0127 |
| 6.75 | 5.562 | 0.4509 | 10 | 5.3476 | 0.9406 | 15 | 0.0321 | 5.5422 | 1.1135 | 10 | 0.0128 | 4.6406 | 1.0893 | 14 | 0.0121 |
| 7 | 5.5006 | 0.3157 | 9 | 5.4268 | 0.9099 | 15 | 0.0053 | 5.5813 | 1.1712 | 11 | 0.0011 | 4.8435 | 0.9954 | 14 | 0.0029 |
| 7.25 | 5.7316 | 0.4677 | 10 | 5.5798 | 0.9922 | 15 | 0.0287 | 5.6999 | 1.305 | 11 | 0.005 | 5.0599 | 1.0898 | 13 | 0.0163 |
| 7.5 | 5.7693 | 0.5011 | 10 | 5.6814 | 1.0814 | 14 | 0.0264 | 5.8517 | 1.2778 | 11 | 0.0095 | 5.1528 | 1.0135 | 13 | 0.0419 |
| 7.75 | 5.6725 | 0.5565 | 9 | 5.7809 | 0.9374 | 15 | 0.142 | 5.9137 | 1.4103 | 11 | 0.0146 | 4.9356 | 1.148 | 14 | 0.0468 |
| 8 | 5.8322 | 0.5159 | 10 | 5.7817 | 0.9535 | 15 | 0.0696 | 5.9205 | 1.3575 | 11 | 0.0076 | 4.9975 | 1.1111 | 14 | 0.0268 |
| 8.25 | 5.9741 | 0.2165 | 9 | 5.8678 | 0.9137 | 15 | 0.0003 | 6.0139 | 1.4328 | 11 | 1E-05 | 4.9845 | 1.302 | 14 | 2E-05 |
| 8.5 | 5.9969 | 0.2922 | 10 | 5.9581 | 0.8392 | 15 | 0.0032 | 5.9972 | 1.3729 | 11 | 8E-05 | 5.2136 | 1.0908 | 12 | 0.0005 |
| 8.75 | 6.0598 | 0.3656 | 10 | 5.9857 | 0.948 | 15 | 0.007 | 5.9425 | 1.3419 | 10 | 0.0006 | 5.2327 | 1.3319 | 14 | 0.0005 |
| 9 | 6.0928 | 0.4458 | 10 | 6.0118 | 0.87 | 14 | 0.0509 | 6.1432 | 1.4449 | 11 | 0.0016 | 5.2926 | 1.4803 | 14 | 0.0011 |
| 9.25 | 6.2512 | 0.3513 | 9 | 6.1603 | 0.8743 | 14 | 0.0145 | 6.3379 | 1.6438 | 10 | 0.0002 | 5.3603 | 1.3848 | 14 | 0.0006 |
| 9.5 | 6.0526 | 0.482 | 9 | 6.2498 | 0.9129 | 15 | 0.0752 | 6.2468 | 1.5358 | 11 | 0.0031 | 5.3843 | 1.3065 | 14 | 0.0083 |
| 9.75 | 6.1074 | 0.395 | 9 | 6.2309 | 1.0106 | 14 | 0.0121 | 6.2177 | 1.4511 | 11 | 0.0011 | 5.2924 | 1.1984 | 14 | 0.0038 |
| 10 | 6.136 | 0.459 | 10 | 6.2361 | 0.8752 | 14 | 0.0589 | 6.1687 | 1.5288 | 11 | 0.0013 | 5.2457 | 1.2564 | 14 | 0.0048 |
| 10.25 | 6.0488 | 0.408 | 9 | 6.3767 | 0.8479 | 15 | 0.0437 | 6.027 | 1.4295 | 10 | 0.0017 | 5.0043 | 1.3672 | 13 | 0.002 |
| 10.5 | 6.2195 | 0.4174 | 10 | 6.4729 | 0.8145 | 15 | 0.0496 | 6.1829 | 1.6737 | 10 | 0.0003 | 5.1054 | 1.4184 | 13 | 0.001 |
| 10.75 | 6.2803 | 0.4517 | 10 | 6.3682 | 0.9055 | 14 | 0.0429 | 6.3209 | 1.6073 | 10 | 0.0008 | 5.1721 | 1.5961 | 13 | 0.0007 |
| 11 | 6.3341 | 0.3559 | 10 | 6.3763 | 0.8877 | 14 | 0.0096 | 6.5214 | 1.5157 | 11 | 0.0002 | 5.1203 | 1.481 | 13 | 0.0002 |

| | | | | | | | | | | | | | | | |
|---|---|---|---|---|---|---|---|---|---|---|---|---|---|---|---|
| 11.25 | 6.4066 | 0.3867 | 10 | 6.4706 | 0.8962 | 15 | 0.0156 | 6.649 | 1.5803 | 11 | 0.0002 | 4.9025 | 1.3317 | 11 | 0.001 |
| 11.5 | 6.4758 | 0.4542 | 10 | 6.5999 | 0.9517 | 15 | 0.0312 | 6.6357 | 1.7403 | 11 | 0.0004 | 5.0771 | 1.7124 | 12 | 0.0004 |
| 11.75 | 6.4485 | 0.5155 | 10 | 6.5739 | 0.909 | 15 | 0.0922 | 6.6975 | 1.8175 | 11 | 0.0008 | 5.1996 | 1.6429 | 12 | 0.0017 |
| 12 | 6.5236 | 0.4084 | 10 | 6.624 | 0.9183 | 15 | 0.0193 | 6.4882 | 1.7596 | 9 | 0.0002 | 5.1414 | 1.5468 | 12 | 0.0004 |
| 12.25 | 6.4793 | 0.4524 | 10 | 6.5784 | 0.9022 | 15 | 0.0431 | 6.4905 | 1.7847 | 10 | 0.0004 | 5.3262 | 1.4507 | 13 | 0.0015 |
| 12.5 | 6.6604 | 0.4404 | 10 | 6.6643 | 0.8497 | 15 | 0.0533 | 6.7684 | 1.7176 | 10 | 0.0004 | 5.3901 | 1.6032 | 13 | 0.0005 |
| 12.75 | 6.5881 | 0.42 | 10 | 6.6356 | 0.9856 | 12 | 0.0162 | 6.5613 | 1.7344 | 11 | 0.0002 | 5.437 | 1.4463 | 13 | 0.0009 |
| 13 | 6.6092 | 0.4359 | 10 | 6.5518 | 0.8575 | 15 | 0.0471 | 6.5275 | 1.67 | 11 | 0.0004 | 5.3784 | 1.4205 | 13 | 0.0013 |
| 13.25 | 6.632 | 0.5269 | 9 | 6.5814 | 0.923 | 13 | 0.1198 | 6.6705 | 1.6756 | 11 | 0.0032 | 5.289 | 1.4146 | 13 | 0.0091 |
| 13.5 | 6.7643 | 0.4158 | 10 | 6.5758 | 0.9361 | 14 | 0.0198 | 6.7727 | 1.8411 | 11 | 0.0001 | 5.378 | 1.3895 | 12 | 0.0012 |
| 13.75 | 6.8556 | 0.3745 | 9 | 6.3009 | 0.7577 | 12 | 0.0555 | 6.9124 | 1.7218 | 11 | 0.0002 | 5.4606 | 1.1804 | 13 | 0.003 |
| 14 | 6.6583 | 0.2913 | 9 | 6.4853 | 0.9072 | 15 | 0.0031 | 6.8332 | 1.6062 | 11 | 6E-05 | 5.569 | 1.1189 | 13 | 0.0007 |
| 14.25 | 6.8064 | 0.4294 | 10 | 6.4691 | 0.8588 | 14 | 0.0436 | 6.561 | 1.8962 | 11 | 0.0001 | 5.5106 | 0.993 | 13 | 0.0171 |
| 14.5 | 6.6335 | 0.4543 | 8 | 6.4034 | 0.8708 | 15 | 0.0903 | 6.7027 | 1.6982 | 11 | 0.0021 | 5.4121 | 1.1067 | 13 | 0.0256 |
| 14.75 | 6.6439 | 0.4573 | 9 | 6.2567 | 0.778 | 12 | 0.1422 | 6.7642 | 1.4246 | 11 | 0.0037 | 5.6579 | 1.2023 | 10 | 0.0122 |
| 15 | 6.7622 | 0.3804 | 9 | 6.4106 | 0.8417 | 13 | 0.0316 | 6.6154 | 1.6312 | 10 | 0.0004 | 5.5081 | 1.043 | 13 | 0.0079 |

**Table S3. Strains used in this study**

| Strain | Genotype | Source |
|---|---|---|
| yJM789 | *Mata SPC110-tdtomato::HIS3 ura3-52 lys2-801 leu2-Δ1 his3-Δ200 trp1-Δ63* | This study |
| yJM2575 | *Matα SPC110-tdTomato::HIS3 ura3-52 lys2-801 leu2-Δ1 his3-Δ200 trp1-Δ63* | This study |
| yJM2578 | *Matα ASE1-GFP::KanMX SPC110-tdTomato::HIS3 ura3-52 lys2-801 leu2-Δ1 his3-Δ200 trp1-Δ63* | This study |
| yJM2579 | *Matα ASE1-GFP::KanMX SPC110-tdTomato::HIS3 ura3-52 lys2-801 leu2-Δ1 his3-Δ200 trp1-Δ63* | This study |
| yJM2580 | *Mata ASE1-GFP::KanMX SPC110-tdTomato::HIS3 ura3-52 lys2-801 leu2-Δ1 his3-Δ200 trp1-Δ63* | This study |
| yJM2581 | *Mata ASE1-GFP::KanMX SPC110-tdTomato::HIS3 ura3-52 lys2-801 leu2-Δ1 his3-Δ200 trp1-Δ63* | This study |
| yJM2756 | *Mata* $ase1^{\Delta 650}$*-GFP::HIS3 SPC110-tdTomato::HIS3 ura3-52 lys2-801 leu2-Δ1 his3-Δ200 trp1-Δ63* | This study |
| yJM2757 | *Mata* $ase1^{\Delta 650}$*-GFP::HIS3 SPC110-tdTomato::HIS3 ura3-52 lys2-801 leu2-Δ1 his3-Δ200 trp1-Δ63* | This study |
| yJM2772 | *Mata* $ase1^{SC}$*-GFP::HIS3 SPC110-tdTomato::HIS3 ura3-52 lys2-801 leu2-Δ1 his3-Δ200 trp1-Δ63* | This study |
| yJM2773 | *Mata* $ase1^{SC}$*-GFP::HIS3 SPC110-tdTomato::HIS3 ura3-52 lys2-801 leu2-Δ1 his3-Δ200 trp1-Δ63* | This study |
| yJM2884 | *Mata* $ase1^{\Delta 693}$*-GFP::HIS3 SPC110-tdTomato::HIS3 ura3-52 lys2-801 leu2-Δ1 his3-Δ200 trp1-Δ63* | This study |
| yJM2885 | *Mata* $ase1^{\Delta 693}$*-GFP::HIS3 SPC110-tdTomato::HIS3 ura3-52 lys2-801 leu2-Δ1 his3-Δ200 trp1-Δ63* | This study |
| yJM3013 | *Mata* $ase1^{3A}$*-GFP::KanMX SPC110-tdTomato::HIS3 ura3-52 lys2-801 leu2-Δ1 his3-Δ200 trp1-Δ63* | This study |
| yJM3053 | *Mata ase1Δ::HIS3 SPC110-tdTomato::HIS3 ura3-52 lys2-801 leu2-Δ1 his3-Δ200 trp1-Δ63* | This study |
| yJM3054 | *Matα ase1Δ::HIS3 SPC110-tdTomato::HIS3 ura3-52 lys2-801 leu2-Δ1 his3-Δ200 trp1-Δ63* | This study |
| yJM3217 | *Mata* $ase1^{3a}$*-GFP::KanMX SPC110-tdTomato::HIS3 TUB1-GFP::LEU2 cdc15-2 ura3-52 lys2-801 leu2-Δ his3-Δ200 trp1-Δ63* | This study |

| | | |
|---|---|---|
| yJM3218 | *Matα ASE1-GFP::KanMX SPC110-tdTomato::HIS3 TUB1-GFP::LEU2 cdc15-2 ura3-52 lys2-801 leu2-Δ1 his3-Δ200 trp1-Δ63* | This study |
| yJM3569 | *Mata ase1$^{Δ693}$-GFP::HIS3 SPC110-tdTomato::HIS3 TUB1-GFP::LEU2 cdc15-2 ura3-52 lys2-801 leu2-Δ1 his3-Δ200 trp1-Δ63* | This study |
| yJM3570 | *Mata ase1$^{Δ693}$-GFP::HIS3 SPC110-tdTomato::HIS3 TUB1-GFP::LEU2 cdc15-2 ura3-52 lys2-801 leu2-Δ1 his3-Δ200 trp1-Δ63* | This study |
| yJM4170 | *Mata ase1$^{3A}$-GFP::KanMX SPC110-tdTomato::HIS3 ura3-52 lys2-801 leu2-Δ1 his3-Δ200 trp1-Δ63* | This study |
| yJM4403 | *Mata trp1-901 leu2-3,112 ura3-52 his3-200 gal4 gal80 Met2::GAL7-lacZ LYS2::GAL1-HIS3 GAL2-ADE2* | James, et al. 1996 |
| yJM4404 | *Matα trp1-901 leu2-3,112 ura3-52 his3-200 gal4 gal80 Met2::GAL7-lacZ LYS2::GAL1-HIS3 GAL2-ADE2* | James, et al. 1996 |
| yJM4408 | *Mata ase1$^{Δ693}$-GFP::HIS3 BIK1-tdTomato::HIS3 ura3-52 lys2-801 leu2-Δ his3-Δ200 trp1-Δ63* | This study |
| yJM4409 | *Mata ase1$^{Δ693}$-GFP::HIS3 BIK1-tdTomato::HIS3 ura3-52 lys2-801 leu2-Δ his3-Δ200 trp1-Δ63* | This study |
| yJM4420 | *Mata ASE1-GFP::KanMX BIK1-tdTomato::HIS3 ura3-52 lys2-801 leu2-Δ his3-Δ200 trp1-Δ63* | This study |
| yJM4421 | *Matα ASE1-GFP::KanMX BIK1-tdTomato::HIS3 ura3-52 lys2-801 leu2-Δ his3-Δ200 trp1-Δ63* | This study |
| yJM4422 | *Mata ASE1-GFP::KanMX BIK1-tdTomato::HIS3 SPC110-mNeonGreen::HIS3 ura3-52 lys2-801 leu2-Δ his3-Δ200 trp1-Δ63* | This study |
| yJM4423 | *Mata ASE1-GFP::KanMX BIK1-tdTomato::HIS3 SPC110-mNeonGreen::HIS3 ura3-52 lys2-801 leu2-Δ his3-Δ200 trp1-Δ63* | This study |
| yJM4434 | *Matα ASE1-GFP::KanMx SPC110-tdTomato::HIS3 cdc15-2 GFP-TUB1::LEU2 ura3-52 lys2-801 leu2-Δ1 his3-Δ200 trp1-Δ63* | This study |
| yJM4436 | *Mata ase1$^{3A}$-GFP::KanMx SPC110-tdTomato::HIS3 cdc15-2 GFP-TUB1::LEU2 ura3-52 lys2-801 leu2-Δ1 his3-Δ200 trp1-Δ63* | This study |
| yJM4591 | *Mata ase1$^{SHNN}$-GFP::KanMX SPC110-tdtomato::HIS3 ura3-52 lys2-801 leu2-Δ1 his3-Δ200 trp1-Δ63* | This study |
| yJM4592 | *Mata ase1$^{SHNN}$-GFP::KanMX SPC110-tdtomato::HIS3 ura3-52 lys2-801 leu2-Δ1 his3-Δ200 trp1-Δ63* | This study |

| | | |
|---|---|---|
| yJM4603 | *Matα ASE1-GFP::KanMX BIM1-tdTomato::HIS3 ura3-52 lys2-801 leu2-Δ1 his3-Δ200 trp1-Δ63* | This study |
| yJM4604 | *Matα ASE1-GFP::KanMX BIM1-tdTomato::HIS3 ura3-52 lys2-801 leu2-Δ1 his3-Δ200 trp1-Δ63* | This study |
| yJM4612 | *Matα ase1$^{\Delta 693}$-GFP::HIS3 BIK1-tdTomato::HIS3 SPC110-mNeonGreen::HIS3 ura3-52 lys2-801 leu2-Δ his3-Δ200 trp1-Δ63* | This study |
| yJM4613 | *Mata ase1$^{\Delta 693}$-GFP::HIS3 BIK1-tdTomato::HIS3 SPC110-mNeonGreen::HIS3 ura3-52 lys2-801 leu2-Δ his3-Δ200 trp1-Δ63* | This study |
| yJM4614 | *Mata ase1Δ::HIS3 BIK1-tdTomato::HIS3 SPC110-mNeonGreen::HIS3 ura3-52 lys2-801 leu2-Δ his3-Δ200 trp1-Δ63* | This study |
| yJM4615 | *Matα ase1Δ::HIS3 BIK1-tdTomato::HIS3 SPC110-mNeonGreen::HIS3 ura3-52 lys2-801 leu2-Δ his3-Δ200 trp1-Δ63* | This study |
| yJM4623 | *Matα ase1$^{SHNN}$-GFP::KanMX BIM1-tdTomato::HIS3 ura3-52 lys2-801 leu2-Δ1 his3-Δ200 trp1-Δ63* | This study |
| yJM4624 | *Matα ase1$^{SHNN}$-GFP::KanMX BIM1-tdTomato::HIS3 ura3-52 lys2-801 leu2-Δ1 his3-Δ200 trp1-Δ63* | This study |

**Table S4. Plasmids used in this study**

| Plasmid | Description | Source |
|---|---|---|
| pJM0014 | *pFA6a-GFP(S65T)-kanMX6* | Petracek and Longtine. 2002 |
| pJM0261 | *pFA6a-mNeonGreen-SpHIS5* | Estrem and Moore. 2019 |
| pSK1050 | *LEU2:GFP-TUB1+3'UTR*, integrating plasmid | Song and Lee. 2001 |
| pJM0049 | *pFA6a-tdTomato::HIS3* | Kurt Thorn |
| pJM0675 | *pDEST-AD2* | Jay Hesselberth |
| pJM0676 | *pDEST-BD2* | Jay Hesselberth |
